# Cytosolic Carboxypeptidase 5 maintains mammalian ependymal multicilia to ensure proper homeostasis and functions of the brain

**DOI:** 10.1101/2024.12.30.630763

**Authors:** Rubina Dad, Yujuan Wang, Chuyu Fang, Yuncan Chen, Yi Zheng, Yuan Zhang, Xinwen Pan, Xinyue Zhang, Emily Swanekamp, Krish Patel, Matthias T. F. Wolf, Zhiguang Yuchi, Xueliang Zhu, Hui-Yuan Wu

## Abstract

Ependymal multicilia position at one-side on the cell surface and beat synchronously across tissue to propel the flow of cerebrospinal fluid. Loss of ependymal cilia often causes hydrocephalus. However, molecules contributing to their maintenance remain yet fully revealed. Cytosolic carboxypeptidase (CCP) family are erasers of polyglutamylation, a conserved posttranslational modification of ciliary-axoneme microtubules. CCPs possess a unique domain (N-domain) N-terminal to their carboxypeptidase (CP) domain with unclear function. Here, we show that a novel mutant mouse of *Agbl5*, the gene encoding CCP5, with deletion of its N-terminus and partial CP domain (designated *Agbl5^M1/M1^*), developed lethal hydrocephalus due to degeneration of ependymal multicilia. Interestingly, multiciliogenesis was not impaired in *Agbl5^M1/M1^* ependyma, but the initially formed multicilia beat in intercellularly diverse directions, indicative of aberrant tissue-level coordination. Moreover, actin networks are severely disrupted and basal body patches are improperly displaced in mutant cells, suggesting impaired cell level polarity. In contrast, *Agbl5* mutants with disruption solely in the CP domain of CCP5 (*Agbl5^M2/M2^*) do not develop hydrocephalus despite increased glutamylation levels in ependymal cilia as similarly seen in *Agbl5^M1/M1^*. This study revealed an unappreciated role of CCP5, particularly its N-domain, in ependymal multicilia stability associated with their polarization and coordination.

## Introduction

Ependymal cells make a single layer of epithelium that lines on the surface of cerebral ventricles. The microtubule-based multicilia formed on the apical surface of ependymal cells beat synchronously to propel the directional flow of cerebrospinal fluid (CSF). Dysfunction of ependymal multicilia is one of major causes of hydrocephalus, which is characterized by abnormal accumulation of CSF in brain ventricle system (Kundishora et al., 2021), often leading to severe neurological disorders such as seizures and mental retardation or even death (Kahle et al., 2016).

Mouse ependymal cells are born from embryonic ventricular zones during brain development and their differentiation initiates in the first postnatal week and completes around P17 (Spassky et al., 2005). Ependymal cells were differentiated from radial glia cells (RGCs) through step-wised procedures, including massive amplification of basal bodies (BBs), docking basal bodies to the apical membrane, multiciliogenesis, polarized positioning and unidirectional alignment of cilia (Lyu et al., 2024; Kyrousi et al., 2017; Ohata and Alvarez-Buylla, 2016). Cytoskeletons microtubule (MT) and filament actin play important yet distinct roles in these procedures. MTs are essential for the assembly, motility, the unidirectional alignment of cilia (namely rotational polarity), and their synchronized beating across the surface of ventricles (Werner et al., 2011; Mirzadeh et al., 2010; Boutin et al., 2014; Arata et al., 2022; Lechtreck et al., 2008; Haycraft et al., 2007; Marszalek et al., 1999), while actin networks are indispensable for BB apical migration, docking (Vladar and Axelrod, 2008), polarized placement (i.e. translational polarity), spacing, and stability (Werner et al., 2011; Mahuzier et al., 2018). However, the molecular clues that contribute to the coordination of these multi-step procedures have not been fully revealed.

Axonemal microtubules undergo a conserved posttranslational modification (PTM)—polyglutamylation, which is processed by tubulin tyrosine ligase like (TTLL) enzymes and forms a polyglutamate side-chain attached to the γ-carboxyl of a glutamate residue in the protein primary sequence (van Dijk et al., 2007; Janke et al., 2005). Conversely, the side-chains can be erased by the 6-member cytosolic carboxypeptidase (CCP) family (Rogowski et al., 2010; Tort et al., 2014). CCP5, encoded by *Agbl5*, is the only identified enzyme that removes the branch point γ-carboxyl linked glutamate after other CCP members shorten the side-chain (Wu et al., 2017; Rogowski et al., 2010). Polyglutamylation has recently been shown to form a specific nano-pattern along axonemal microtubules to regulate ciliary beating (AlvarezViar, et al., 2024). Consistently, deregulation of axonemal glutamylation in mice results in phenotypes related to primary ciliary dyskinesia (Yang et al., 2021; Ikegami et al., 2010).

Although the prototypic CCP member Nna1/CCP1 has been primarily linked to neurodegeneration (Rogowski et al., 2010), accumulating evidence including that from analysis of the evolutionary history of CCP expressing organisms suggests the cilium-and basal body-related role of this family (Rodriguez de la Vega Otazo et al., 2013; Wang et al., 2023; He et al., 2018; Rodriguez-Calado et al., 2023). In zebrafish, knocking down CCP5 induces ciliopathic phenotypes, such as body curvature and hydrocephalus, suggesting an important role of CCP5 in ciliary motility and function (Lyons et al., 2013; Xie et al., 2022; Zhang et al., 2018). Surprisingly, *Agbl5* knockout (KO) mice with targeted disruption of the carboxypeptidase (CP) domain in CCP5 do not exhibit overt anomalies other than male infertility due to defective tail formation of developing sperm (Wu et al., 2017; Giordano et al., 2019), raising the question of its importance in mammalian epithelial motile cilia.

In addition to the CP domain, CCPs share an N-terminal domain (ND, so called N-domain) unique for this family that is folded into an antiparallel β sandwich anchored to the CP domain and contributes to the recognition of substrates (Kalinina et al., 2007; Rimsa et al., 2014; Chen et al., 2024). We and other groups reveal that CCP5 and CCP6 interact with centriolar proteins in ciliated cells (Wang et al., 2023; Rodriguez-Calado et al., 2023). In particular, CCP5 interacts with CP110 through its ND (Wang et al., 2023), indicating its role as a protein-interacting interface. It is thus possible that the previously reported *Agbl5*-deficient mice do not carry a null mutant due to their potential expression of the N-domain.

To clarify the physiological significance of CCP5, we generated a novel *Agbl5* KO allele (designated *Agbl5^M1^*), in which the deletion was extended to the N-terminus of CCP5. Strikingly, *Agbl5^M1/M1^*mice developed severe hydrocephalus and died before the age of 2 months. Interestingly, the multicilia of *Agbl5^M1/M1^* ependymal cells initially formed but underwent serious degenerations. Moreover, the initially formed multicilia in the mutant ependyma exhibit diverse intercellular beating directions and their BB patches are not properly displaced. Therefore, CCP5 is involved in intracellular positioning and intercellular coordination of mammalian ependymal multicilia that are critical for their maintenance to ensure proper homeostasis and functions of the brain.

## Results

### Mice carrying a null *Agbl5* mutant (*Agbl5^M1/M1^*) display markedly reduced lifespan

*Agbl5* KO mouse models with targeted deletion of a region encoding its carboxypeptidase domain did not exhibit overt anomaly other than male infertility (Wu et al., 2017; Giordano et al., 2019). In order to determine whether the N-domain of CCP5 possesses additional biological function, we generated a novel *Agbl5* KO allele (designated as *Agbl5^M1^*) using CRISPR-CAS9 genome manipulation system. In this allele, the region from the start codon to intron 8, which encodes the sequence including the N-domain and part of CP domain, was replaced with a tdTomato reporter (Figure 1A). The possible off-targets were excluded by examining the top 10 possible sites predicted for each single guide RNA (sgRNA) in descendants of 2 independent founders (Table S1). The absence of *Agbl5* transcripts in the brain, eye, spinal cord, spleen, and testis of *Agbl5^M1M1^* animals was confirmed by RT-PCR using primers targeting the region encoding the CP domain (Figure 1B).

**Figure 1.**
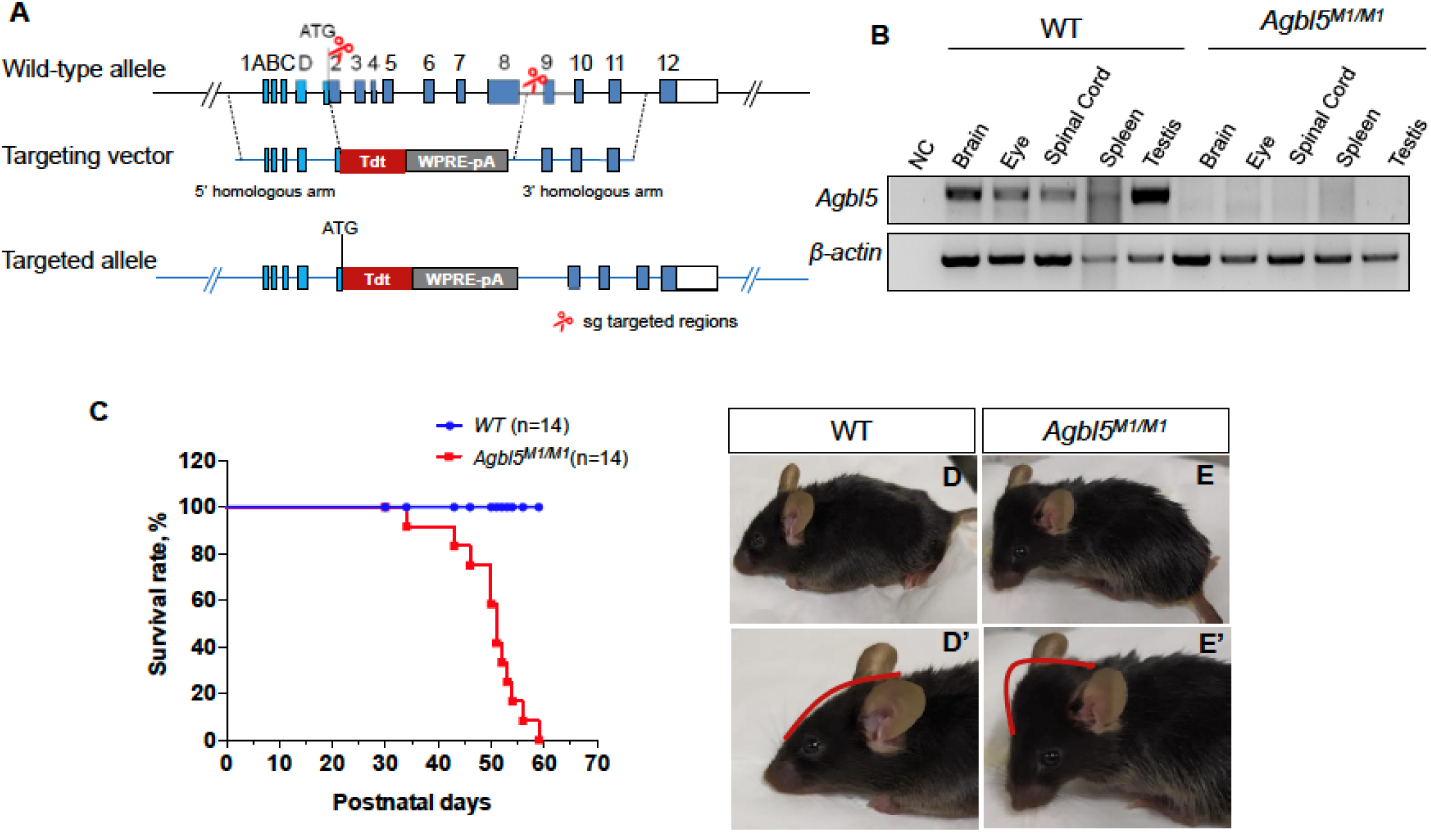
A novel *Agbl5* mutant mouse model exhibited domed head and reduced lifespan. (A) Schematic representation of the knock-out/knock-in strategy to create the novel *Agbl5* mutant (*Agbl5^M1^*) allele. A region between start codon and intron 8 was replaced with a tdTomato reporter cassette using CRISPR/CAS9 system. (B) RT-PCR using primers targeting a region spanning exon 6 and exon 8 of *Agbl5* confirmed the absence of *Agbl5* transcripts in brain, eye, spinal cord, spleen, and testis of *Agbl5^M1/M1^* mice. NC, negative control. (C) The Kaplan-Meier survival curve showed that the *Agbl5^M1/M1^* mice hardly survived more than 50 days (*p*<0.0005, log-rank (Mantel-Cox) test). (D-E’) Representative images of P50 wild-type (D, D’) and *Agbl5^M1/M1^* (E, E’) mice. Different from the wild-type mice (D, D’), the mutant (E, E’) developed dome-shaped head as indicated with red lines (D’, E’).

Pregnant *Agbl5^M1^* heterozygous females do not exhibit obvious anomaly. The *Agbl5^M1/M1^* pups appeared normal at the birth. Compared with the wild-type animals, however, *Agbl5^M1/M1^* mice showed reduced body weight between P17-P21, and stopped growth after P30 (Figure S1). At the age between P41 and P51, the homozygous mutant mice even gradually lost the weight (Figure S1). Surprisingly, unlike the previously reported *Agbl5* KO models (Wu et al., 2017; Giordano et al., 2019), *Agbl5^M1/M1^* mice hardly survive longer than 54 days (Figure 1C).

### *Agbl5^M1/M1^* mice developed severe hydrocephalus

Upon visual inspection, we noticed that the *Agbl5^M1/M1^*mice developed dome-shaped cranium around P40 (Figure 1D-E’) with a full penetration. When brains from mutants about P50 were dissected, we found that they were filled with fluid— a characteristic feature of hydrocephalus (data not shown).

Cerebrospinal fluid (CSF) is produced from choroid plexuses and fills the ventricles of the brain. It flows from the lateral ventricles (LV), through the interventricular foramens, and into the third ventricle, cerebral aqueduct, and the fourth ventricle. To assess the alteration in ventricles of *Agbl5^M1/M1^*mice, serial coronal sections along the axis of the brain were microscopically examined after hematoxylin-eosin staining. Compared with those of wild-type mice (Figure 2A-C), the lateral ventricles (Figure 2D), both the dorsal and ventral 3^rd^ ventricles (Figure 2D, D’), and the fourth ventricle (Figure 2F) of the *Agbl5^M1/M1^*mutant mice were dramatically enlarged, while the aqueduct was altered to a less extent (Figure 2B, E). These results indicate that this mutation in *Agbl5* likely affects a common process required for all ventricles of brain.

**Figure 2.**
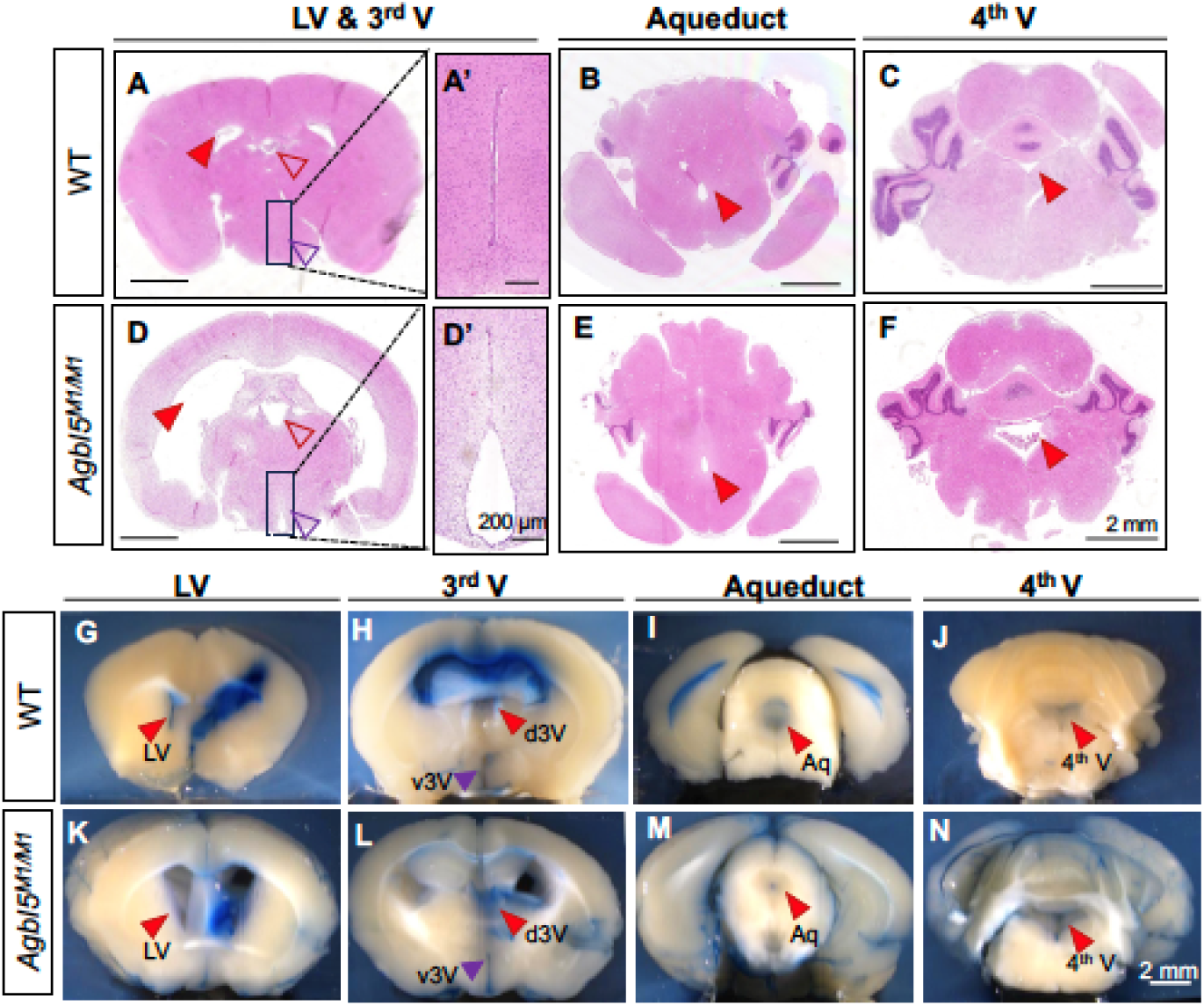
Histological and flow assessment of the cerebrospinal fluid (CSF) pathway. Hematoxylin-Eosin staining on a serial coronal brain sections of P46 wild-type (WT, A-C) and *Agbl5^M1/M1^* (D-F) mice. Markedly enlarged lateral ventricles (LV, D, closed arrowhead), the third ventricles (D, open arrowheads, D’), and the fourth ventricle (F) were observed in *Agbl5^M1/M1^* mice compared to the corresponding part in wild-type mice (A, B and C, arrowheads), while the size of the aqueduct was less affected in the mutant (B vs. E, arrowheads). A’ and D’ are the close view of ventral 3^rd^ ventricles that were indicated in dashed boxes in A and D respectively. The dorsal and ventral 3^rd^ ventricles are pointed with open arrowheads in red and purple respectively in A and D. (G-N) Flow assessment of CSF in mice of P30 intraventricularly injected with Evans Blue. Whole brains were fixed in 4% PFA 20 min later after injection to allow ink distribution and diffusion in the brain of wild-type (G-J) and mutant mice (K-N). Tissue sections showed that diffused ink was clearly observed in the lateral ventricles (LV, red arrowheads in G, K), the third dorsal (d3V, red arrowheads in H, L) and ventral ventricles (v3V, purple arrowheads in H, L), the fourth ventricles (4^th^V, red arrowheads in J, N) and aqueducts (Aq, red arrowheads in I, M) of both wild-type (G-J) and mutant (K-N) mice, indicative of communicating hydrocephalus. Scale bars: 2 mm for A-N; 200 μm for A’, D’.

We further assessed whether the hydrocephalus was caused by the blockage of the CSF flow path. The dye Evans Blue was injected into the right lateral ventricle of P30 wild-type and *Agbl5^M1/M1^* mice. An obstruction in CSF path would delay the appearance of ink in the other ventricular compartments (Zou et al., 2020). We found that for both wild-type and mutant mice, the injected dye diffused into the left lateral (Figure 2G, K), the third (Figure 2H, L), and the fourth ventricles (Figure 2J, N), suggesting a communicating hydrocephalus instead a blockage of CSF flow path in *Agbl5^M1/M1^* mice.

### Ependymal multicilia in *Agbl5^M1/M1^* hydrocephalic mice are defective in both number and motility

The flow of CSF is propelled by the movement of multicilia of ependymal cells that line on the surface of ventricles. To determine the cause of hydrocephalus in *Agbl5^M1/M1^*mice, the whole-mount lateral walls of lateral ventricles from P45 mice were immunostained for acetylated tubulin (Ac-tub), which is abundantly present in ciliary axoneme (Delgehyr et al., 2015). In the wild-type mice, the bundles of Ac-tub positive multicilia evenly covered the surface of ventricle walls and their tips pointed to the same direction (Figure 3A). In contrast, Ac-tub positive multicilia bundles were _sp_arsely detected on the ventricle walls of the mutant mice, and often with the unbundled cilia lying on the cell surface (Figure 3B). Scanning electron microscopy (SEM) analysis of lateral ventricles of P30 mice further revealed that unlike the ventricle walls of wild-type mice, which are covered with unidirectional multicilia bundles (Figure 3C, E), multicilia on the ventricle walls of mutant mice were severely lost and often with only a few cilia left in the middle of the cell surface (Figure 3D, F). These results point to the defects of multicilia in *Agbl5^M1/M1^*ependymal cells.

**Figure 3.**
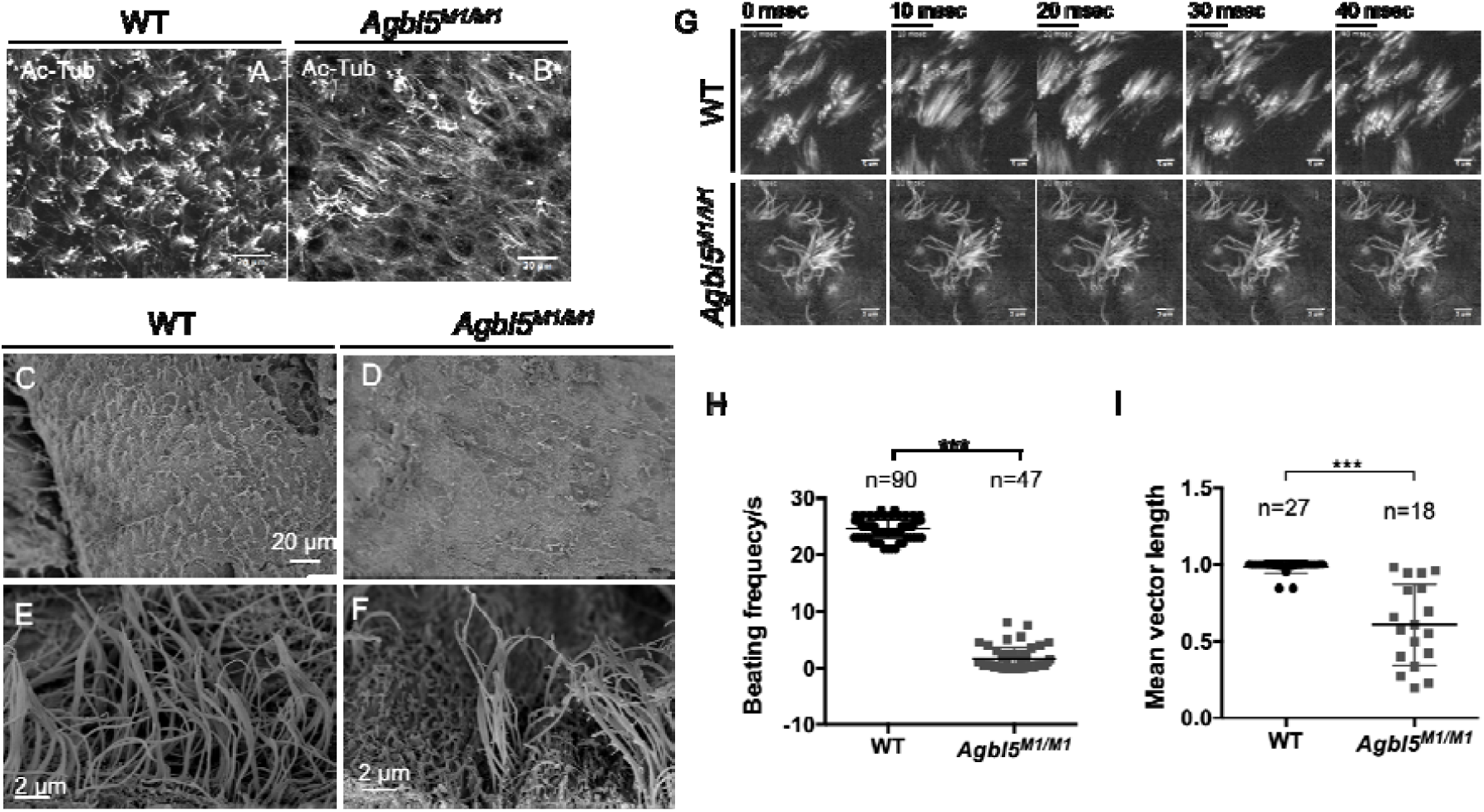
Aberrant ependymal multicilia in *Agbl5^M1/M1^*hydrocephalic mice. (A-B) Representative images of the whole-mount lateral walls of LVs from P45 wild-type (WT, A) or *Agbl5^M1/M1^* (B) immunostained with the ciliary marker, acetylated tubulin (Ac-Tub). While multicilia in WT ependyma evenly cover the ventricle surface and point to the same direction, mutlicilia bundles were scattered with many Act-Tub positive cilia lying on the cell surface. (C-F) Scanning electron microscopy analysis of LV walls from wild-type (C, E) and *Agbl5^M1/M1^* (D, F) mice of P30. In the wild-type LV, ependymal cell were covered with evenly distributed cilia bundles in a uniformed direction (C), while in the mutant mice, cilia only in the middle of ependymal cell surface remain (D). (E, F) Higher magnification image showed that the length of remaining ependymal cilia in *Agbl5^M1/M1^* mice (F) is similar to that in wild-type animal (E). (G) Sequential images of ciliary beating in wild-type (upper row) and *Agbl5^M1/M1^* (lower row) mice. (H-I) Quantification of beating frequency of ependymal cilia (H) or the consistency of their beating directions (reflected by mean vector length) in individual cells (I) of P45 mice (from 3 wild-type and 2 mutant mice). Scale bar, A-D, 20 μm; E-F, 2μm, G, 5 μm.

It is possible that the loss of ependymal multicilia is secondary to other causes in *Agbl5^M1/M1^* mice. To this end, we assessed the beating patterns of the ependymal multicilia in adult mice using high-speed imaging technique. In the wild-type mice, ependymal multicilia beat unidirectionally in a synchronized pattern (Figure 3G, Movie 1). In *Agbl5 ^M1/M1^*ependymal cells, however, the majority of remnant multicilia hardly move (Figure 3G, Movie 2), while the others beat much slower than those in wild-type (Figure 3H, Movie3). Those cilia still mobile in the mutant largely remain a normal stroke pattern, but the beating directions of individual cilia in the same cluster are often diverse (Movie 3, quantified in Figure 3I). These observations suggested a defective rotational polarity. Taken together, in *Agbl5^M1/M1^*, the hydrocephalus is indeed directly associated with the dysfunction of ependymal cilia.

To determine whether loss of CCP5 also affects multicilia in other organs, we assessed trachea multicilia in *Agbl5^M1/M1^* using SEM. At the age of P30, the multicilia of tracheal epithelial cells in wild-type orientate in same directions (Figure S2A, B). However, in tracheas of the mutant, multicilia in the same cells often radiate to different directions, yet the number appears less than that in the wild-type (Figure S2C, D). Therefore, the function of CCP5 is commonly required for alignment and maintenance of multicilia.

### Basal bodies are largely dispersed in ependymal cells of *Agbl5^M1/M1^* hydrocephalic mice

The actin networks in ependymal cells not only contribute to the polarized placement of basal bodies (BB), but are also important to maintain their stability against the sheering force of CSF (Werner et al., 2011; Mahuzier et al., 2018; Hirota et al., 2010). We wondered whether the positioning of BBs and the integrity of actin networks were also affected in *Agbl5^M1/M1^*ependyma. Whole-mount lateral walls of the lateral ventricles (LVs) were co-stained for F-actin and CEP164, a distal appendage protein of BB (Graser et al., 2007). We found that unlike the wild-type ependymal cells where BBs were clustered at one side of cell surface and well organized in rows (Figure 4B-C, E, F, Figure S3A), about 30% of *Agbl5^M1/M1^* ependymal cells contain dispersed BBs (Figure 4H-I, K, L, Figure S3B). Even in the cells with clustered BBs, the BBs were often not properly aligned (Figure 4I, K, Figure S3B), suggesting possible defects on subapical actin network that is required for BB spacing (Mahuzier et al., 2018). Notably, in the mutant ependyma, BBs were not detected in a substantial portion of cells (Figure 4L).

**Figure 4.**
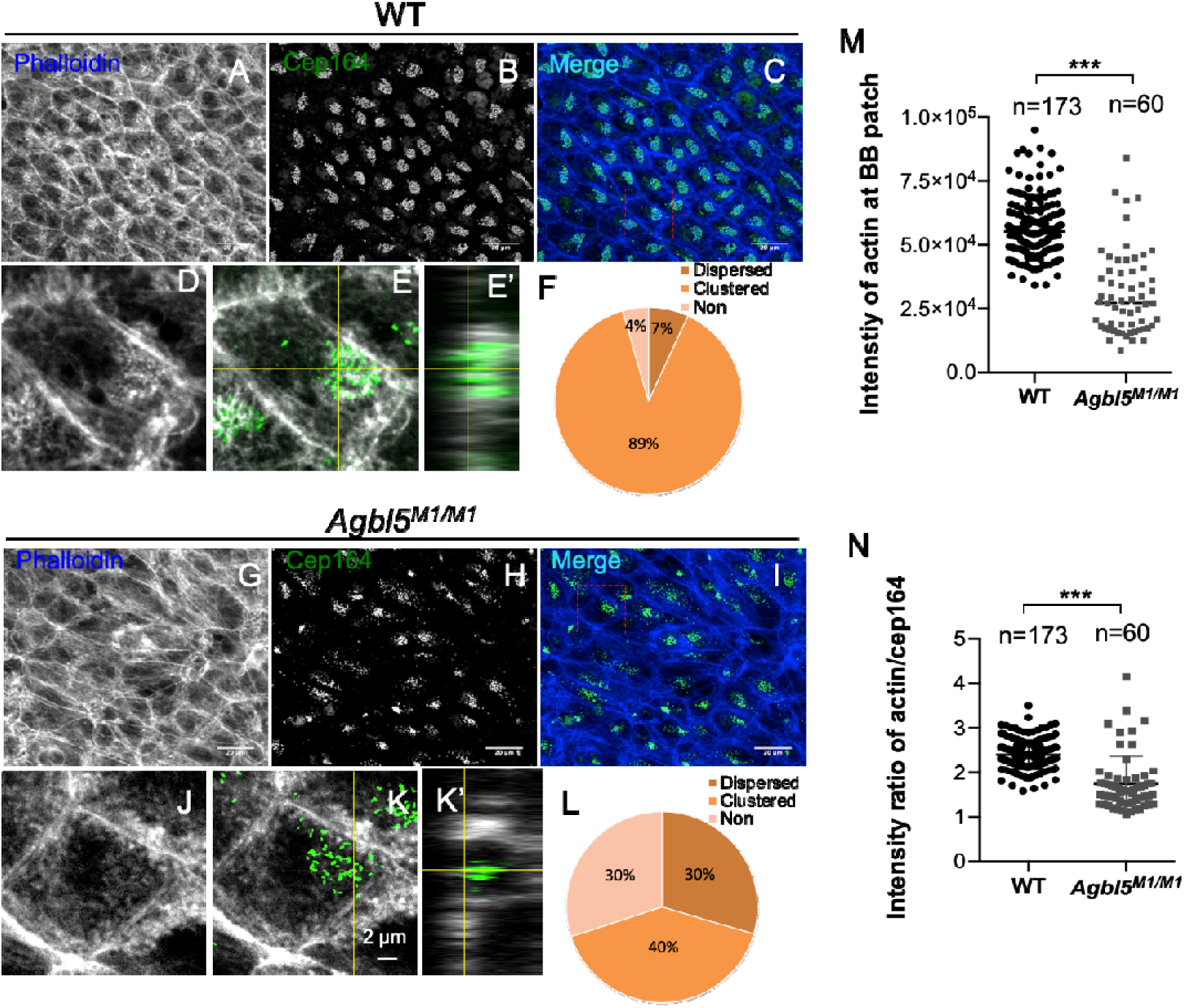
The apical actin network in *Agbl5^M1/M1^* ependymal cells is disrupted. (A-L) immunostained with the centriolar distal appendage marker CEP164 (B, H) with actin network labeled with phalloidin (A, G, G, J). While BBs are clustered and polarized in the wild-type ependymal cells (B, C), those in the mutant (H, I) are often diffused. The actin networks are largely disrupted in the mutant ependymal cells G, J, vs. A, D for wild-type). (D-E’, J-K’) Z-projection views of apical actin network around BB in wild-type (C, E) and *Agbl5^M1/M1^* (J, K) ependymal cells and respective orthogonal views (E’, K’). The mutant ependymal cells lack the compact actin networks even around clustered BBs. (F, L) Quantification of ependymal cells with differently distributed BBs in wild-type (F) and *Agbl5^M1/M1^* mice (L). (M, N) Quantification of the total intensity of F-actin around BBs (M) and the intensity of F-actin per BB (N). Scale bar, A-C, G-I-, 20 μm; D-E’, J-K’2 μm.

Consistent with the observation of abnormal BB positioning in *Agbl5^M1/M1^*ependymal cells, we found that their apical actin networks, which contribute to polarization and stabilization of BBs (Mahuzier et al., 2018), were severely disrupted (Figure 4G, I, J vs 4A, C, D for wild-type). Z-project images further revealed that in ependymal cells of wild-type mice, a compact apical actin patch is polarized to one side of the cell and colocalized with CEP164 immunosignals (Figure 4D-E’). In contrast, this specialized actin patch was abolished in *Agbl5^M1/M1^* ependymal cells even around BBs that are clustered (Figure 4J-K’). Quantitative analysis showed that the intensity of actin patch around BBs in the mutant ependyma is significantly reduced compared with that in wild-type (Figure 4M). This is not attributed to the variation in BB number between two genotypes, as the intensity of F-actin per BB in the mutant is also significantly lower than that in the wild-type (Figure 4N). Therefore, in *Agbl5^M1/M1^* ependymal cells, the actin networks are affected, which possibly impairs stability of BBs and their multicilia against the sheering of CSF.

### *Agbl5* is expressed in ependymal cells

Comparing the size of lateral ventricles in the wild-type and mutant mice at different postnatal stages, we found that the enlarged ventricles consistently appeared since P7 in the mutant mice (Figure S4A). Taking the advantage of the tdTomato reporter constructed in the *Agbl5^M1^*allele, we assessed the spatial expression of *Agbl5* in the heterozygous mice. At P7, tdTomato signal was detectable in ependymal cells and broadly expressed in other regions of the brain, but much lower in the cells in the subventricular zone (Figure S4B, C). At P12, tdTomato signals were highly expressed in the ependymal cell layer of all walls of LV (Figure. 5A, B), but were still largely devoid from the subventricular zone (Figure 5B, B’, arrow head). In the dorsal-lateral niche of the ventricle wall, tdTomato positive cells were not restricted to the single layer on the ventricle surface, but expended a few layers laterally (Figure 5B’, arrow). Co-immunostaining of brain sections for tdTomato and S100β, a marker of mature ependymal cells (Jacquet et al., 2009; Spassky et al., 2005; Ohata et al., 2014) further confirmed the identify of tdTomato-positive cells along the ventricle surface in *Agbl5^WT/M1^* mice (Figure 5C-E). This spatial expression pattern supports a possible role of *Agbl5* in ependymal cell development.

**Figure 5.**
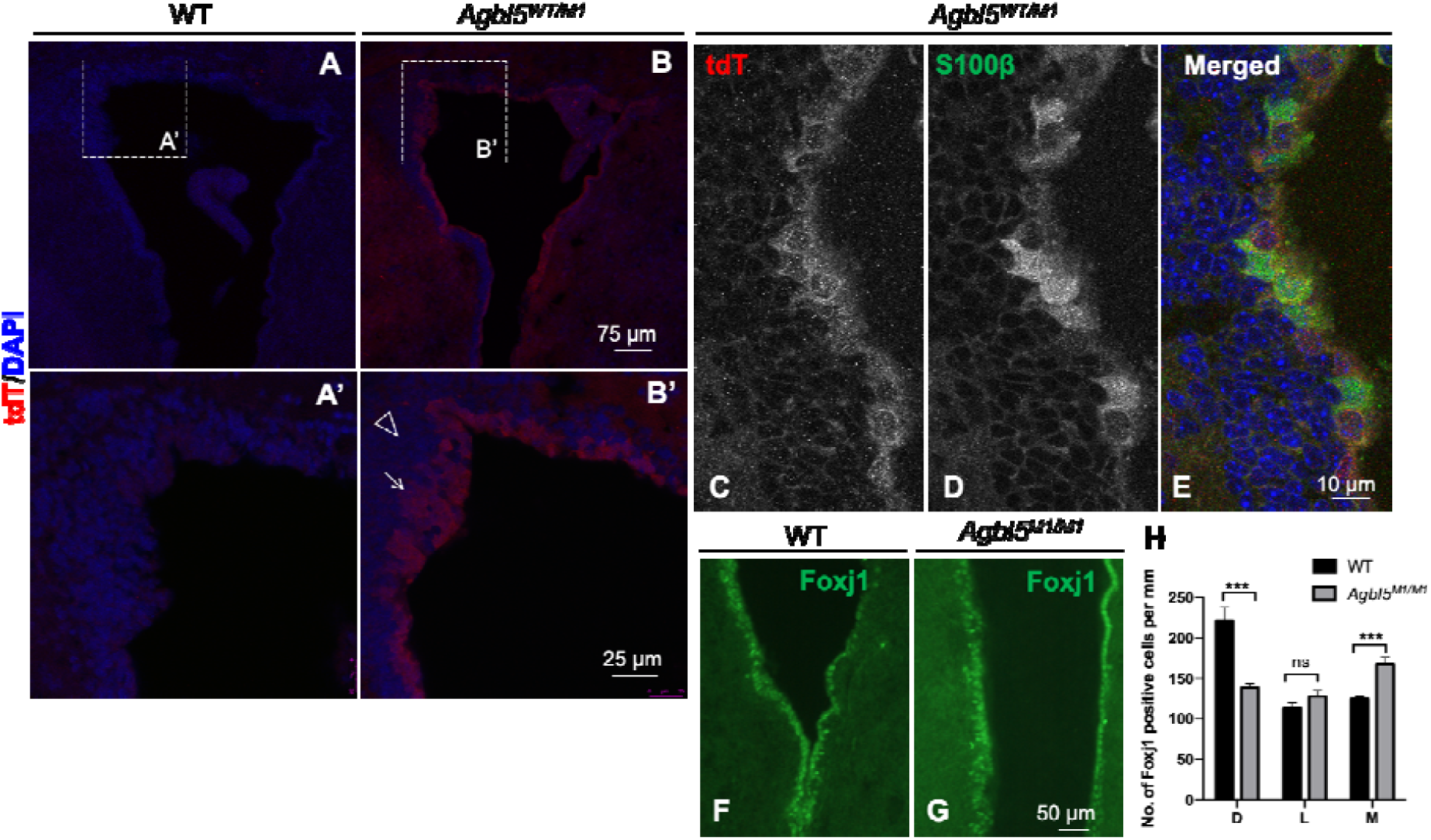
Expression of multiciliogenesis-promoting protein is not impaired in *Agbl5^M1/M1^* ependyma. (A-B’) Immunofluorescence analysis revealed that tdTomato signals can be detected in heterozygous *Agbl5^M1^* (*Agbl5^WT/M1^*) brain (B, B’), but not in the wild-type control (A, A’). The tdTomato signals were localized in the ependymal cells but largely devoid from the subventricular zone (arrowhead). At the dorsal-lateral region of the LV, the tdTomato signals extend to 2-3 layers (arrow). (C-E) In brain section of P12 *Agbl5^WT/M1^* mice, tdTomato expression is colocalized with that of S100β, an ependymal cell marker along the surface of lateral ventricles. (F, G) Lateral ventricles from P7 wild-type and *Agbl5^M1/M1^* mic were immunostained with Foxj1, a protein promoting multiciliation. (H) Quantification showed that th number of Foxj1-positive cells per length of LV walls in the mutant mice (n=5) is decreased in the dorsal wall, but unchanged or slightly increased in the lateral and middle walls respectively compared to that in the wild-type mice (n=5). Error bars represent SEM. ***, *p*<0.001; student’s *t*-test. Scale bars, A, B, 75 μm; A’, B’, 25 μm; C-E,10 μm, F-G, 50 μm.

Ependymal cells are born in the embryonic stages and differentiated from mono-ciliated radial glia cells (RGCs) through multiple step-wise procedures, including cell fate commitment, multiciliogenesis, and maturation (Marques et al., 2019). Their maturation predominantly takes place in the first 2 postnatal weeks (Spassky et al., 2005). We wondered whether the commitment of RGCs to the fate of ependymal cells in *Agbl5^M1/M1^* mice was affected. The expression of vimentin, a protein that marks ependymal cells since embryonic stages (Schnitzer et al., 1981; Vidovic et al., 2018) was assessed in P7 mice. It was found that cells lining along the ventricle walls of both wild-type and mutant mice were similarly positive for vimentin staining (Figure S5), suggesting that in *Agbl5^M1/M1^* mice, the commitment of RGCs to ependymal cell fate was not altered.

Foxj1 is a transcription factor that promotes multiciliation in ependymal cells (Jacquet et al., 2009). We next determined whether the expression of Foxj1 is altered in *Agbl5^M1/M1^* mice. At P7, the number of Foxj1-positive cells along each wall of lateral ventricles in *Agbl5^M1/M1^* mutants was increased compared to that of the wild-type mice (Figure S6), but the number of Foxj1-positive cells per length unit is significantly reduced in the dorsal wall probably due to the greatest elongation of this wall. In other two walls, the number of Foxj1-positive cells is similar or slightly increased in the mutant (Figure 5F-I). These results suggested that the reduction of multicilia in *Agbl5^M1/M1^* ependymal cells was not caused by insufficient expression of Foxj1.

### *Agbl5* loss leads to increased glutamylation level in the lateral ventricle

Among the 6 CCP family members, CCP5 is the only one catalyzing the removal of branch point glutamate in the modified proteins (Rogowski et al., 2010; Wu et al., 2017; Kimura et al., 2010). The glutamylation levels in *Agbl5^M1/M1^*lateral ventricle were assessed using GT335 antibody, which recognizes the branch point glutamate (Wolff et al., 1992), and polyE antibody, which detects more than 3 glutamate residues at C-termini respectively (Figure 6A) (Rogowski et al., 2010; Lacroix et al., 2010). Similar to the other parts of the brain (Rogowski et al., 2010), tubulin glutamylation level in the lateral ventricles is low at P7, but is greatly increased at P21 and remains at a similar level at P30 (Figure 6B). Compared with wild-type mice, loss of *Agbl5* led to an increased GT335 level in the lateral ventricles of all ages examined, but did not change the immunoreactivity for polyE (Figure 6B). These results are similar to what was observed in cerebella of previously reported *Agbl5* KO mice (Wu et al., 2017), reflecting a phenomenon related to CCP5 enzyme activity.

**Figure 6.**
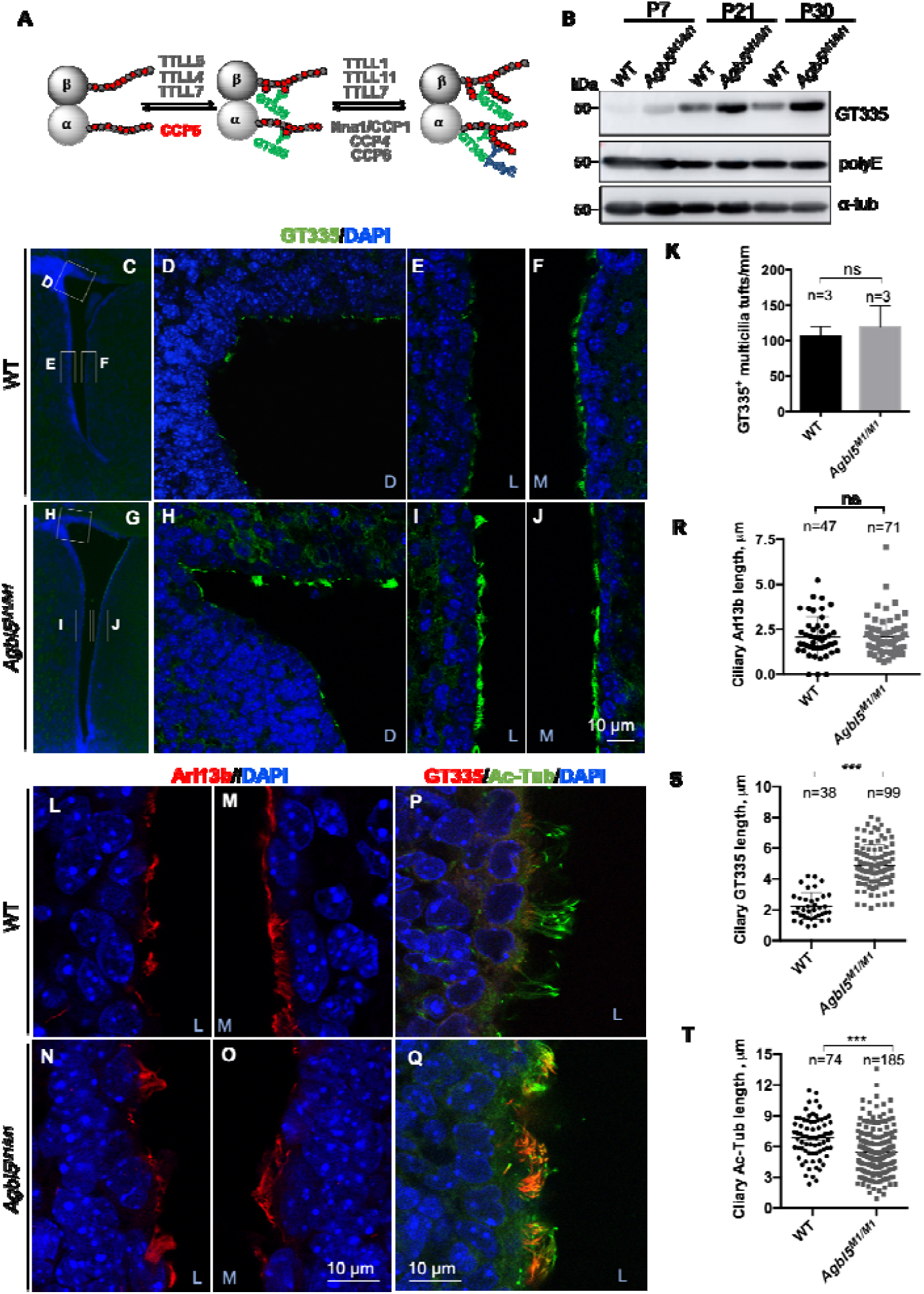
The glutamylation level is increased in ependymal multicilia of *Agbl5^M1/M1^* mice. (A) A schematic representation shows the enzymes involved in tubulin polyglutamylation and modification recognized by GT335 and polyE antibodies respectively. (B) Immunoblotting of LV from mice of different ages showed that compared to the wild-type, the immunosignals of GT335 but not that of polyE are increased in *Agbl5^M1/M1^* mice at all ages examined. (C-J) Lateral ventricles of P7 wild-type (C-F) and *Agbl5^M1/M1^* (G-J) mice stained with GT335 (green) and DAPI. Representative images show that the intensity and length of GT335 immunosignals in ependymal cilia are increased in all three (H, dorsal; I, lateral; J, middle) walls of LV in the mutant mice compared with the respective walls in wild-type LVs (D, dorsal; E, lateral; F, middle). (K) The number of multicilia tufts is comparable between the wild-type (n=3) and mutant mice (n=3) in the lateral walls. (L-O) LVs of wild-type (L, M) and *Agbl5^M1/M1^* (N, O) mice immunostained for Arl13b (red) with nuclei visualized by DAPI staining. (R) Quantification showed that compared to that of the wild-type mice (n=3), the length of Arl13b signal in ependymal multicilia of *Agbl5^M1/M1^*mice (n=3) were not changed. (P-Q) LVs of wild-type (P) and *Agbl5^M1/M1^*(Q) mice co-immunostained for GT335 (red) and acetylated tubulin (Ac-Tub, green). (S,T) Quantification showed that compared to that of the wild-type mice (n=3), the length of ciliary GT335 signal (S) in ependymal multicilia of *Agbl5^M1/M1^* mice (n=3) is significantly increased while that of Ac-Tub (T) is reduced. Letters in blue: D, dorsal wall; L, lateral wall, M, middle wall. Error bars represent SEM, student’s *t*-test. Scale bars, C, G, 100 μm; D-J, L-Q, 10 μm.

As microtubules in multicilia are glutamylated, we assessed whether loss of *Agbl5* affected the glutamylation level in ependymal multicilia using immunofluorescence (IF). Ependymal cells differentiate during the first postnatal week (Spassky et al., 2005). At P7, the GT335-positive multicilia appear as short tufts in the wild-type LV (Figure 6C-F), with only a few in the dorsal wall, but more in the lateral and middle walls (Figure 6C-F). In the *Agbl5^M1/M1^* mice, the number of GT335-positive multicilia tufts was comparable with that of wild-type mice (Figure 6K), but the intensity (Figure 6D-F vs. 6H-J) and the length of GT335 signal in the multicilia (Figure 6R) were both strongly increased. Therefore, loss of *Agbl5* indeed increased the glutamylation level in ependymal multicilia.

To determine whether increased glutamylation signals in the cilia leads to increased length of cilia, we assessed the signal of Arl13b, a protein localized on ciliary membrane (Caspary et al., 2007). We found that the length of the ciliary Arl13b signals was not significantly different between wild-type and mutant mice (Figure 6K-N, Q), indicating that loss of *Agbl5* did not alter the length of ependymal cilia at this stage.

To determine whether loss of *Agbl5* affects other types of tubulin PTM in ependymal multicilia, we co-stained brain sections for glutamylation and tubulin acetylation (Ac-Tub), another conserved PTM of ciliary axoneme. In wild-type mice, GT335 immunosignals are mainly located at the base of ependymal multicilia, while signals of Ac-Tub extend in the axoneme (Figure 6O, P). In the mutants, however, GT335 signals stretched far into the axoneme (Figure 6O, P) and can reach the length of Ac-Tub signals in some cases. Interestingly, compared with that in wild-type mice (Figure 6O-P, S), the length of acetylated tubulin signal in the multicilia of mutant mice were reduced (Figure 6O-P, S). Therefore, loss of *Agbl5* increased the level of tubulin glutamylation at the expense of acetylation level in multicilia, reflecting a crosstalk between two types of modifications.

### Ependymal cilia in *Agbl5^M1/M1^* are initially motile

To determine whether the ependymal multicilia initially formed in *Agbl5^M1/M1^* mice function properly, we recorded their beating using a high-speed camera in P15 mice. At this stage, ependymal multicilia of *Agbl5^M1/M1^*were largely preserved (Figure 7A, B) and were motile (Figure 7C, Movie 5). Those bundled multicilia beat slightly faster than that of wild-type mice (Figure 7D). Moreover, in contrast to wild-type ependyma where multicilia of neighboring cells beat in similar directions (Figure 7A, C, E, Movie 4), those in mutant mice often beat in different directions (Figure 7B, C, E, Movie 5). In some individual ependymal cells of the mutant mouse, cilia in the same cell beat in different directions (Figure 7C, closed arrowheads; Movie 5, arrows). Therefore, the multicilia in *Agbl5^M1/M1^* ependymal cells are initially formed, but their intra-and intercellular beating coordination are impaired.

**Figure 7.**
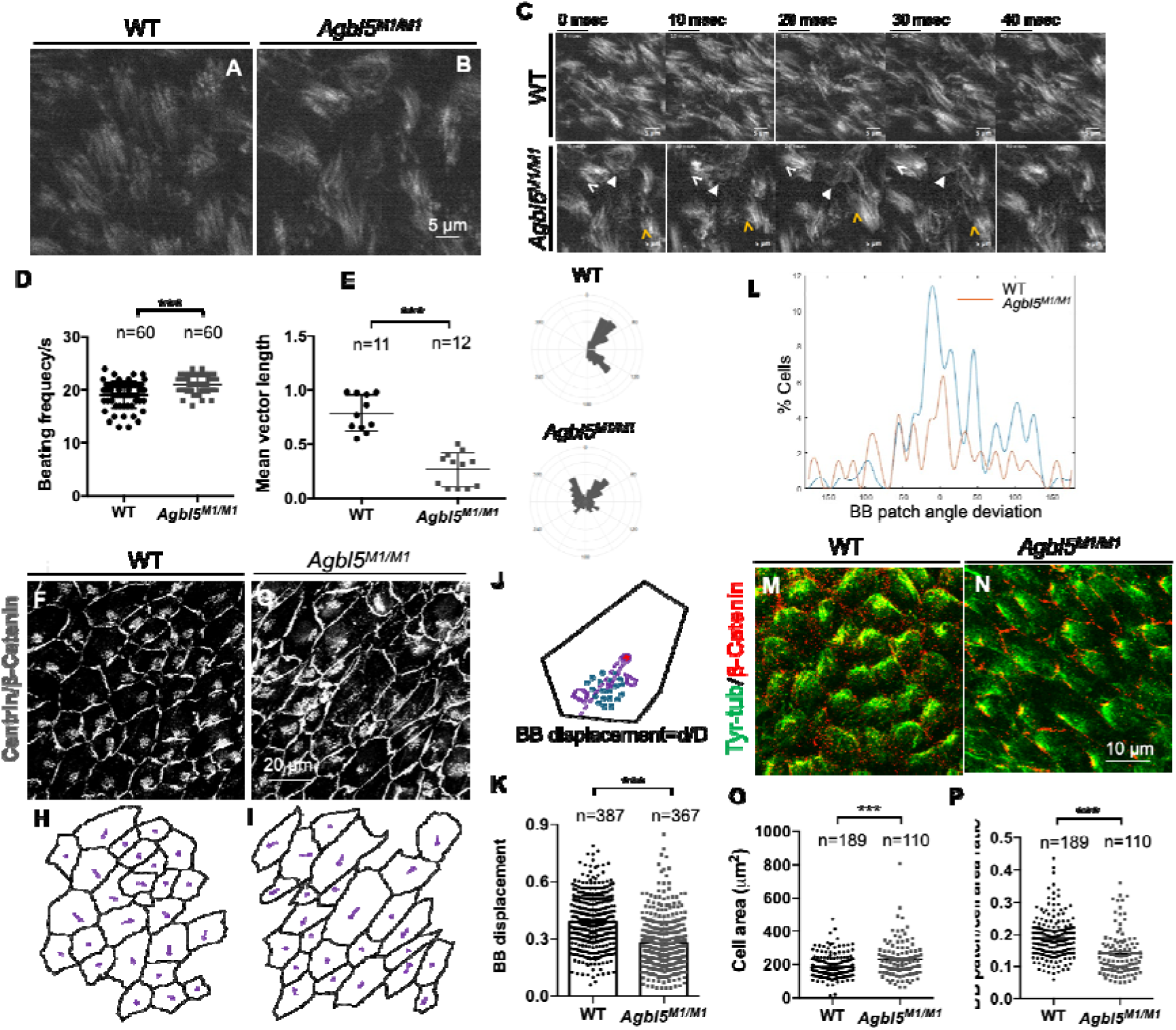
The initially formed ependymal multicilia in *Agbl5^M1/M1^* mice are motile. (A-B) Images of SiR-tubulin labeled whole-mount LVs from P15 wild-type (A) and *Agbl5^M1/M1^* (B) mice show that ependymal multicilia are initially formed in the mutant. (C) Sequential images of ciliary beating of P15 wild-type (upper row) and *Agbl5^M1/M1^* (lower row) showed that the multicilia of wild-type ependymal cells beat in similar direction, while that of mutant are asynchronously. White and yellow open arrowheads indicate respective beating directions of multicilia of two cells; the closed arrowhead points to multicilia of an individual cell beat in opposite directions. (D) Bundled *Agbl5^M1/M1^* multicilia largely beat at the frequency slightly higher than that of wild-type (n=30 for each animal, 2 mice for each genotype). Error bars represent SD. (E) The consistency of cilia beating directions of WT and *Agbl5^M1/M1^* ependymal cell in the tissue level reflected by the mean vector length of individual imaging field. Representative histograms of beating angles for each genotype, represented in polar coordinates. The area of each wedge is proportional to the percentage of angles in the corresponding angle range. (F, G) Whole-mount LVs from P15 wild-type (F) and *Agbl5^M1/M1^* (G) mice were co-immunostained with Centrin (BB marker) and β-Catenin (cell boundary marker). (H, I) Traces of the intercellular junction labeled with β-Catenin of ependymal cells shown in F and G respectively. The purple arrows show the vectors drawn from the center of the apical surface to that of the BB patch. (J) Diagram showing the measurement of BB patch displacement. (K) Quantification showed that BB patches in *Agbl5^M1/M1^* ependymal cells are not properly displaced (n= 387 from 5 wild-type mice; n=367 from 4 mutants), *p*<0.001, student’s *t* test. (L) Histogram of the distribution of BB patch angles in ependymal cells of WT (blue) and *Agbl5^M1/M1^* (orange), (n=327 from 5 wild-type; n=233 from 4 mutants), *p*<0.001, Watson’s 2-sample U^2^ test. (M, N) Whole-mount lateral walls of LV from P10 WT (M) and *Agbl5^M1/M1^*(N) co-immunostained for β-Catenin and tyrosinated tubulin (Try-tub) showed that the polarization of MTs was not affected in ependymal cells of the mutant. (O, P) Quantification of ependymal cell apical surface delineated by β-Catenin staining showed that the mutant (N) exhibits larger area than that of wild-type (M), while the ratio of the area of BB patch over the area of apical surface is reduced in the mutant (n=189 from 4 wild-type and n=110 from 3 mutants). Scale bars, A-C, 5 μm; F, G, 20 μm.

### The displacement of BB patches in *Agbl5^M1/M1^* ependymal cells is impaired

In P45 *Agbl5^M1/M1^* mice, BBs are dispersed over the surface of ependymal cells (Figure 4H, I, K). It is possible that the aberrant BB positioning at this late stage is secondary to the loss of multicilia, because ependymal ciliary beating induces the formation of actin networks that stabilize BBs (Mahuzier et al., 2018). To this end, we assessed the BB displacement in ependymal cells of mice at P15, when the ependymal multicilia are still present in the mutant. The whole-mount of the lateral walls of LVs were co-immunostained with Centrin and β-catenin to label the BBs and cell boundary respectively (Figure 7F, G). The displacement of BB patches is measured as the distance between the center of BB patch and that of the apical cell surface normalized to the distance between the cell surface center and boundary in the same direction (Figure 7J, (Ohata et al., 2014; Mirzadeh et al., 2010)). We found that the displacement of BB patches in *Agbl5^M1/M1^*ependymal cells is significantly reduced compared with that in wild-type (Figure 7K).

In differentiated ependymal cells, BB patches tend to align in similar directions among the neighboring cells, reflecting a tissue-level polarity (Mirzadeh et al., 2010; Hirota et al., 2010; Ohata et al., 2014; Takagishi et al., 2017). We measured the BB angles by drawing vectors from the center of cell apical surface to the center of the BB patch in each cell (Figure 7H, I). The distribution of BB patch angles is significantly more diverse in the mutant than that in the wild-type ependyma (Figure 7L). These results suggest that *Agbl5* contributes to BB positioning at early stage.

### The microtubule network coordinating neighboring cells are polarized in *Agbl5^M1/M1^* ependyma

At the apical surface of multi-ciliated cells, two sets of microtubule networks are involved in coordination of ciliary beating. In addition to one set that goes along with the actin meshwork and underlines the patch of BBs, another set is polarized and extended between the patch and the cell cortex, providing an anchoring point at the cell cortex in the similar location in neighboring cells (Boutin et al., 2014). The latter plays a role in the tissue-level coordination of multicilated cells. Previous studies showed that microtubule +TIPs and tyrosinated tubulin, which marks newly synthesized microtubule-plus ends, are polarized to the anchoring point of the cell cortex (Takagishi:2017bn; Vladar et al., 2012). Given the uncoordinated ciliary beating amongst neighboring cells in *Agbl5^M1/M1^* ependyma and the nature of tubulin modification enzyme of CCP5, we assessed the expression of tyrosinated tubulin (Try-Tub) at P9 when the density of tyrosinated tubulin reaches highest on the anterior side (Takagishi et al., 2017). We found that similar to wild-type mice, tyrosinated tubulin is polarized to the anterior side in ependymal cells of the mutant and anchors to the cell cortex (Figure 7M, N). Therefore, loss of CCP5 in *Agbl5^M1/M1^* mice did not alter tubulin polarization that coordinates neighboring ependymal cells.

The MT network around basal bodies coordinates neighboring cilia in the same cell (Werner et al., 2011). We wondered whether it is glutamylated in ependymal cells and whether loss of CCP5 increased its glutamylation levels. We analyzed confocal images of whole-mount LV walls co-stained with GT335 and Centrin antibodies. GT335-labeled axoneme of multicilia in *Agbl5^M1/M1^* is longer and less organized compared to that of wild-type (Figure S7A, B, E, F). The GT335 signals under the BB layer (localized by Centrin immunosignals) are much weaker than those in the cilia and appear denser in the area round BBs (Figure S7C, D, G, H). However, the GT335 signals in this layer were not obvious different between two genotypes (Figure S7C, D, G, H). Therefore, the glutamylation level of MT network around basal bodies is not affected by *Agbl5^M1/M1^* ependymal cells.

### The apical surfaces of *Agbl5^M1/M1^* ependymal cells are expanded

Primary cilia of radial glia cells (RGCs), the progenitors of ependymal cells, preset signals for their translational polarity (Mirzadeh et al., 2010; Ohata et al., 2015). CCP5 is localized in primary cilia (He et al., 2018; Wang et al., 2023). Previous studies showed that abolishing primary cilia in RGCs, results in diverse orientations of BB patches with reduced displacement (Mirzadeh et al., 2010), similar to what is seen in *Agbl5^M1/M1^* ependymal cells. Loss of primary cilia in RGCs also leads to expanded apical surfaces and tightly clustered BB patches (Mirzadeh et al., 2010). Similarly, in *Agbl5^M1/M1^* ependyma, the apical cell surfaces are increased and the area of BB patch accounts for 14.1±0.6% of the apical surface, compared to 18.8±0.5% in the wild-type (Figure 7 O, P). However, in differentiating ependymal cells of *Agbl5^M1/M1^* mice, primary cilia are present, with a length comparable with that is the wild-type (Figure S8). Therefore, the aberrant translational polarity of ependymal cells in *Agbl5^M1/M1^* may also be associated with signals transduced by RGC primary cilia.

### Loss of the enzyme activity of CCP5 alone is not sufficient to cause hydrocephalus

In order to confirm that the loss of ependymal cilia in *Agbl5^M1^*mutants did not solely result from the inactivity of CCP5 enzyme, we generated a second *Agbl5* mutant allele (designated *Agbl5^M2^*) where a region spanning exon 6 and intron 8 that encodes a part of CP domain of CCP5, was replaced with IRES-guided tdTomato reporter (Figure 8A). This mutant mimics the one we reported previously (Wu et al., 2017). The absence of *Agbl5* transcripts was confirmed in the testis, brain, and eye of *Agbl5^M2/M2^* mice (Figure 8B).

**Figure 8.**
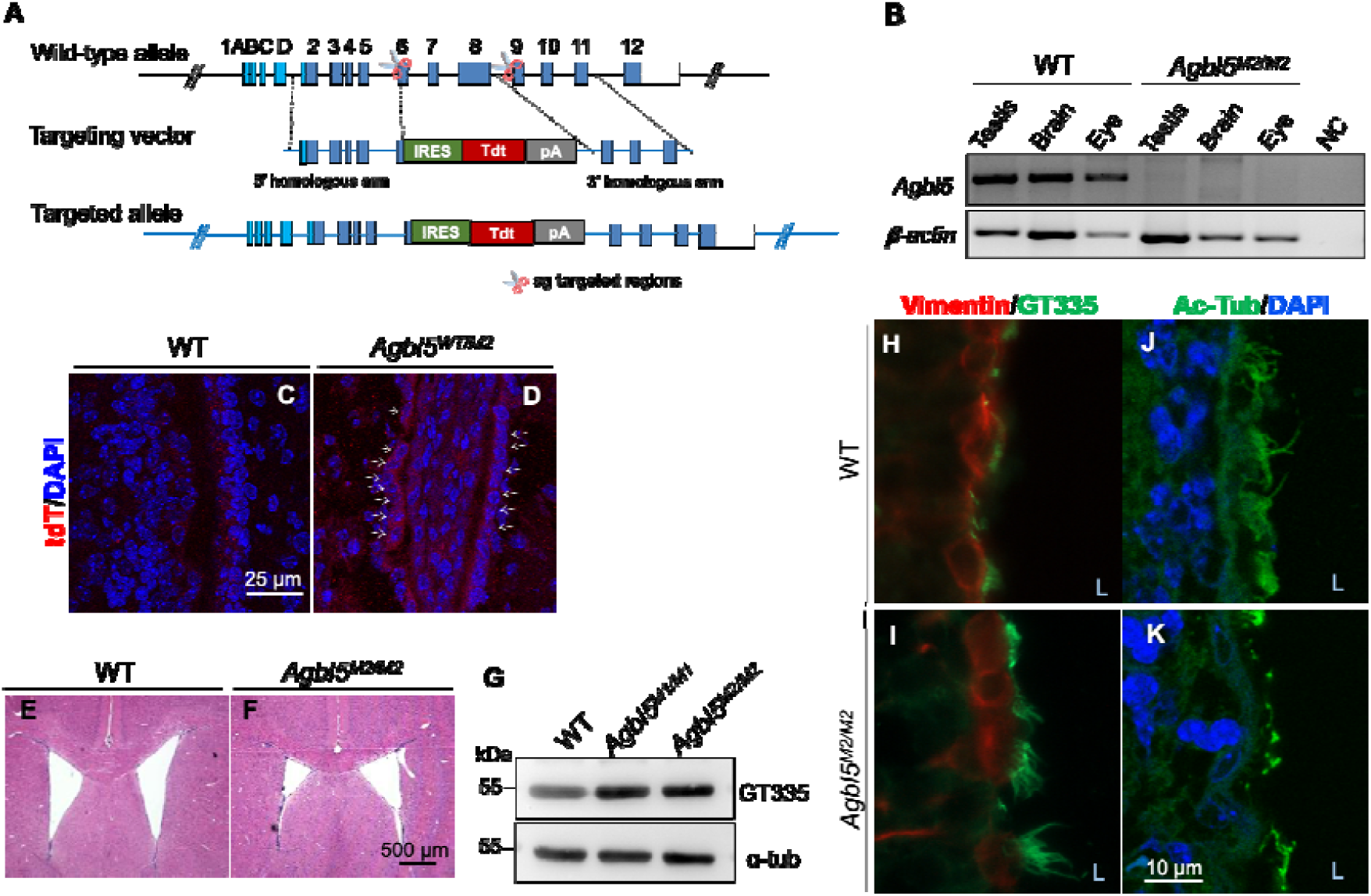
Targeted disruption of CP domain alone in *Agbl5* did not cause hydrocephalus, despite the increased glutamylation in ependymal cilia. (A) Schematic representation of the knock-out/knock-in strategy to create a second *Agbl5* mutant (*Agbl5^M2^*) allele that resembles the one used in previous studies (Wu et al., 2017). (B) RT-PCR using primers targeting deleted region in *Agbl5^M2^* allele confirmed the absence of *Agbl5* transcripts in brain, eye, and testis in *Agbl5^M2/M2^* mice. NC, negative control. (C-D) Similar to that in *Agbl5^WT/M1^* mice, tdTomato immunosignal is detected in ependymal cells of P7 *Agbl^M2^* heterogenous mice (D, arrows). (E-F) Hematoxylin-Eosin staining of coronal sections of brains from 3-month old wild-type (E) and *Agbl5^M2/M2^* (F) mice, where no enlarged ventricles were observed. (G) Immunoblotting assay showed that the tubulin glutamylation level was increased in the brain of both *Agbl5* mutants compared to that in the wild-type. (H-I) Immunostaining showed that the ciliary GT335 signals in ependymal cells of *Agbl5^M2/M2^* mice (I) are increased compared with that of the wild-type (H). (J-K) The ciliary acetylated-tubulin (Ac-Tub) signals are reduced in ependymal cilia in *Agbl5^M2/M2^* mic (K) compared with that in the wild-type (J). L, lateral wall. Scale bars, C, D, 25 μm; E, F, 500 μm; H-K, 10 μm.

Traced with the tdTomato reporter, the expression of *Agbl5* was also detected in the ependymal cell layer of the lateral ventricles in *Agbl5^M2^*heterozygous mice at P7 (Figure 8C, D), similar to our observations in the *Agbl5^M1^* allele (Figure 5B, Figure S4C). Consistent with previously reported *Agbl5* KO mice (Wu et al., 2017; Giordano et al., 2019), *Agbl5^M2/M2^* do not exhibit obvious anomaly other than male infertility. No enlarged cerebral ventricles were observed even in 3-month-old mice (Figure 8E, F).

We wondered whether mutations in two *Agbl5* mutants alter the glutamylation level to different degrees. Immunoblotting assessment revealed that compared with the wild-type, the GT335 immunoreactivity was increased in *Agbl5^M2/M2^* brain to an extent similar to that in *Agbl5^M1/M1^* mice (Figure 8G). Immunofluorescence assay further revealed that similar to that in *Agbl5^M1/M1^* mice, the intensity of GT335 signal in ependymal multicilia was also increased, accompanied by the reduction in the immunosignals of acetylated tubulin (Figure 8H-K). These results emphasized that loss of CCP5 enzyme activity contributes to the altered PTM levels in ependymal multicilia of both *Agbl5* mutants, which though is not necessary to be deleterious.

Therefore, the elevated glutamylation level in *Agbl5* mutants are attributed to the loss of CCP5 enzyme activity, which alone is not sufficient to induce hydrocephalus.

## Discussion

Polyglutamylation is a conserved PTM on cilia axonemal microtubules. In contrary to its writers that have been linked to ciliary architecture and motility, the role of polyglutamylation erasers in cilia has not been fully appreciated. This is largely due to the absence of a broad spectrum of ciliopathic phenotypes (other than photoreceptor degeneration and male infertility) in their mammalian mutants. In this study, using a null *Agbl5* allele (*Agbl5^M1^*), we demonstrated that *Agbl5* is essential for the development of ependymal cells in cerebral ventricles to maintain brain homeostasis. Particularly, deletion of the N-terminal domain of CCP5 along with part of its CP domain impairs the stability of multicilia in ependymal cells, leading to lethal hydrocephalus. In this mutant, ependymal multicilia are initially formed but the synchronousness of their beating is compromised. Moreover, the BB patches in individual cells are not properly positioned and the compact apical actin networks around basal bodies are disrupted. In contrast, the mutation solely targeting the CP domain of CCP5 did not cause enlarged ventricles, despite the altered PTM status in ependymal cilia. This study uncovers a novel role of *Agbl5* and emphasizes the requirement of its N-domain in ependymal multicilia development.

Individual ependymal cells exhibit two types of planar cell polarity (PCP), i.e. rotational polarity (the unidirectional orientation of cilia) and translational polarity (the off-centered positioning of BB patch on the cell surface). These two types of PCP extend at tissue level, reflected by coordinated cilia orientation and positioning across tissue (Mirzadeh et al., 2010; Mahuzier et al., 2018; Boutin et al., 2014; Arata et al., 2022). In *Agbl5^M1/M1^* mice, the multicilia are initially formed, but their intra-and inter-cellular beating coordination is compromised, suggesting an aberrant rotational polarity. In addition, the displacement of BB patches in individual ependymal cells is also impaired, indicating a defect in translational polarity. Therefore, CCP5 contributes to the establishment of both rotational and translational polarities in ependymal cells.

It is unexpected that loss of CCP5, a MT modification enzyme exhibits profound effects on actin-related functions in ependymal cells, at least in two aspects. Firstly, loss of CCP5 in *Agbl5^M1/M1^* impairs the displacement of BB patches in ependymal cells. The establishment of translational polarity in ependymal cells relies on the actomyosin dynamics (Hirota et al., 2010). Particularly, the phosphorylation of non-muscle myosin 2 (NMII) is essential for the polarized localization of BB patches (Hirota et al., 2010). It will be intriguing to further determine how the expression and/or localization of phosphorylated NMII are altered in *Agbl5^M1/M1^*ependymal cells. Secondly, the apical actin networks in the mutant ependymal cells are severely disrupted in *Agbl5^M1/M1^* ependymal cells, while the BBs in ependymal cells are not properly aligned, suggesting a possible defect in the sub-apical actin networks around BBs as well (Werner et al., 2011; Mahuzier et al., 2018; Arata et al., 2022). It remains unknown how CCP5 is involved in assembly of actin networks in ependymal cells. A recent study demonstrated that the regulators of tubulin polyglutamylase complex are involved in MT association of actin nucleators and MT–actin crosslinkers, exemplified the role of polyglutamylation enzymes in bridging functions between these two types of cytoskeleton (Wang et al., 2022). Given the important role of apical actin networks in maintaining the stability of BBs (Mahuzier et al., 2018), its aberrance likely contributes to the loss of ependymal multicilia in *Agbl5^M1/M1^*in response to the sheering of CSF. Moreover, the impaired local synchronization of cilia can cause the steric collision (Ringers et al., 2023), which may generate additional force to stub the cilia rooted in a uncompact actin network.

Planar cell polarity (PCP) pathway (i.e. non-canonical Wnt pathway) interplays with MT dynamics to establish the rotational polarity in ependymal cells, but only fine-tunes the translation polarity (Ohata et al., 2014; Boutin et al., 2014). In mature ependymal cells, a set of MTs that is polarized between the BB patch and the cell cortex provides contributes to synchronize the beating of multicilia across the tissue (Boutin et al., 2014). Although the glutamylation level is robustly increased in cilia of *Agbl5^M1/M1^* ependymal cells, it is barely affected in MT around BBs or where else in cytoplasm, suggesting a spatially restrict regulation of CCP5 activity. The polarized MT network, detected with tyrosinated-tubulin, is thought to depend on the asymmetric localization of the transmembrane PCP protein Fzd (Boutin et al., 2014; Vladar et al., 2012). As the polarization of this set of MT in *Agbl5^M1/M1^* is not affected, the localization of Fzd assumptively remain normal. It is tempting to conceive that CCP5 may directly regulate PCP components. A recent study revealed that protein Disheveleds (Dvls), the cytosol adaptors of PCP pathway are polyglutamylated at their C-termini, which regulates the distribution of these molecules in the condensates (Kravec et al., 2024). However, CCP5 does not degrade the α-linked polyglutamate at the C-termini of Dvls despite their colocalization. It requires further investigation how CCP5, particularly its N-domain is involved in PCP pathway to establish the rotational polarity in ependymal cells.

In monociliated cells, we showed that CCP5 interacts with the ciliation negative regulator CP110 through its N-domain (Wang et al., 2023). Although loss of CP110 also caused asynchronous ciliary beating, reduced motility and randomization of directionality, or a complete loss of motility in multicilia cells (Walentek et al., 2016), similar to what were seen in *Agbl5^M1/M1^* ependymal cells, we did not detect significant reduction of CP110 expression *in Agbl5^M1/M1^* lateral ventricles (Figure S9). Different from primary cilia, CP110 localizes to the motile cilia-forming basal bodies. In multiciliated cells, it is specially needed for ciliary adhesion complex formation (Antoniades et al., 2014) and basal body interaction with the actin cytoskeleton (Walentek et al., 2016). It requires further investigation how CCP5 and CP110 coordinate in multicilia development or whether CCP5 requires additional mediators to facilitate the unique polarized placement of multicilia in ependymal cells.

The earliest clue of the translational polarity of ependymal cells resides in the asymmetric positioning of primary cilia in their progenitors, RGCs. Other than the presence of primary cilia per se, activation of the sensory molecules localized in cilia of RGCs, such as Pkd1 and Pkd2 are critical to properly polarize RGCs and thereby ependymal cells (Ohata et al., 2015). Notably, knocking down CCP5 increased the ciliary targeting of Pkd2 in cell models (He et al., 2018). Further assessment of the positioning of primary cilia and the ciliary targetting of Pkd2 or other proteins in *Agbl5^M1/M1^* RGCs may shed light on the underpinnings of defective PCP in their ependymal cells.

Using two *Agbl5* mutant models, this study revealed the role of CCP5 beyond its enzyme activity. It was surprising that although loss of CCP5 enzyme activity results in elevated glutamylation level in ependymal cilia, it is not necessary to be deleterious. Prominently, loss of CCP5 increased glutamylation level at the expense of tubulin acetylation in ependymal cilia. Similar observation was reported in the connecting cilia of retina in an *Agbl5* mutant mouse (Aljammal et al., 2024). Although these two modifications take place at different locations in the MT, both are characteristic of stable MTs. Therefore, a complementary/compensation mechanism that regulates the balance between these two modifications in axoneme apparently exists.

Supported by *in vivo* evidence, the present study reinforces the speculation that the unique N-domain in CCP family may have a role specifically related to cilium/centrioles (Wang et al., 2023; Rodriguez de la Vega Otazo et al., 2013). In *Agbl5^M2/M2^* mice, the transcripts coding CCP5 N-terminus are still present (data not shown), making it possible that this region still fulfills the function to regulate or recruit other proteins and thereby mitigates the deleterious effects (Wu et al., 2017; Giordano et al., 2019). Such candidates may include other CCP members (e.g. CCP6) that degrade long chain polyglutamate of non-tubulin substrates important for cilium/centrioles function (Hao et al., 2021; Kravec et al., 2024). Moreover, multicilia in young A*gbl5^M1/M1^*mice beat in a frequency slightly higher than that in the wild-type, but at later stages the remanent cilia are either paralyzed or beat much slower, raising the question how loss CCP5 interferes with the machinery of cilia beating. Although polyglutamylation directly regulates axonemal microtubule-dynein interaction in motile cilia, this modification does not affect axonemal structure (Kubo:2010da; Suryavanshi:2010br; Alvarez Viar et al., 2024). In addition, the ependymal multicilia in *Agbl5^M2/M2^* remain intact in adult, thus excluding the possibility that the reduced beating velocity of cilia in adult *Agbl5^M1/M1^* ependyma is caused by the increased glutamylation level. It is possible that loss of CCP5-ND compromises the function or ciliary targeting of certain proteins required for maintaining the integrity of cilia structure during aging, leading to impaired motility. Given the unique possession of the N-domain in CCP family, additional investigation focusing on the interaction partners and function of this domain can further provide insights in this regard.

Taken together, this study uncovered an unappreciated function of *Agbl5* in ependymal cell development, and provides a novel clue contributing to the positioning, coordination, and maintenance of ependymal multicilia mediated by a polyglutamylation enzyme though beyond its role to modify the ciliary axoneme.

## Materials and Methods

### Animals

Mice colonies were maintained in a SPF animal facility on a 12-hr light: 12-hr dark cycle with free access to food and water. All animal study protocols were approved by the Institutional Animal Care and Use Committee (IACUC) at Tianjin University.

### Generation of novel *Agbl5* mutant alleles

Two *Agbl5* mutant alleles (*Agbl5^M1^* and *Agbl5^M2^*) were created using CRISPR-CAS9 gene editing system on C57BL/6N background by Biocytogen Co., Beijing, China. To generate *Agbl5^M1^* allele, sgRNA 5’-AGCAGCAGACCCACTAGCGG-3’ and 5’-TCTCAGCTCTATGGAAGACG-3’, which target the sequence next to the start codon and intron 8 respectively, together with the donor plasmid harboring tdTomato reporter cassette flanked with 3’-and 5’-homologous arms, and Cas9 mRNA were injected into zygotes with well-recognized pronuclei. To generate *Agbl5^M2^* allele, sgRNA 5’-AACATGAAAGCCGTATTCTTGGG-3’ and 5’-TCTATGGAAGACGGGGGTGCTGG-3’, which target exon 6 and intron 8 respectively, together with the donor plasmid harboring an IRES guided tdTomato reporter cassette flanked with 3’- and 5’- homologous arms, and Cas9 mRNA were injected into zygotes with well-recognized pronuclei. The founders (F0) were determined by genotyping PCR. After bred with wild-type mice, the F1 mice with desired mutation were selected after genotyping, and the integration and copy number of the inserted fragment were further validated with Southern blotting. *Agbl5* mutant heterozygous mice were inbred to produce homozygous and wild-type litter mates. The *Agbl5* mutant and wild-type alleles were genotyped using primers listed in Table S3.

### Off-target determination

The online CRISPR-CAS9 target prediction tool CCTOP (Stemmer et al., 2015) or COSMID (Cradick et al., 2014) were used to predict off-targets of the sgRNAs used in generation of *Agbl5* mutant alleles. Eleven top-ranking predicted off-targets for each sgRNA were selected and primers to amplify the predicted genomic region were designed using primer3 software. With the genome DNA from heterozygous offspring of 2 independent founders as the template, the amplicons of predicted off-targets were obtained using *pfu* polymerase enzyme and subsequently subjected to Sanger sequencing after gel extraction. Sequences of amplified predicted off-target regions of wild-type and mutants were aligned to the sequences from UCSC genome browser using CLCBIO main workbench software. The absence of predicted off-targets was confirmed in both *Agbl5* mutant alleles. The assessed list of predicted off-targets were provided in Tables S1 and S2, and the sequencing results are available upon request.

### RT-PCR analysis of *Agbl5* expression

Total RNA was extracted from testis, brain, eye, and spleen of P45 *Agbl5* mutant mice or wild-type litter mates using Trizol reagent (CWBIO, Taizhou, China) according to manufacturer’s protocol. First strand cDNA was synthesized using ABScript first strand cDNA synthesis kit (ABclonal, Wuhan, China) according to manufacturer’s instructions. Expression of *Agbl5* was analyzed by subsequent PCR using primers 5’-TCTCTGGATGGACTTCGTGT-3’ and 5’-TGGTTCGTGGGACTCTTGG-3’. *β-actin* amplified using primers 5’- ATATCGCTGCGCTGGTCGTC-3’ and 5’-AGGATGGCGTGAGGGAGAGC-3’ was used as a control.

### Detection of tdTomato fluorescence

Mice were subjected to transcardiac perfusion with PBS followed by 2% paraformaldehyde (PFA) after anesthesia with 10% chloral hydrate. Brains were dissected and fixed with 2% PFA overnight at 4^°^C. Brains were embedded in Tissue-Tek OCT (Sakura, China) after cryoprotection in 30% sucrose and sectioned in 20 μm thick coronally. After wash with PBS, brain sections were subjected to confocal images directly.

### Analysis of the flow of CSF using Evans Blue

The circulation of CSF was analyzed by using Evans Blue dye as previously described (Liu et al., 2016). Mice of age P30 were anesthetized with Avertin. The 30-gauge needle attached to syringe was positioned at specific bregma points of 0.1 mm posterior and 1.0 mm on the head. Five microliters of Evans blue dye (diluted as 4% in PBS) were carefully and gradually injected into the right lateral ventricle (LV) of each mouse. 20 min later, mice were sacrificed and whole brains were dissected and fixed in 4% PFA for 12 hr. The samples were then embedded in agarose (4% in PBS) and kept at 4°C for 30 mints to solidify. Coronal sections were cut to inspect the presence of dye in each part of the ventricular system. Images were captured by using a stereomicroscope (SMZ1270, Nikon, Japan) equipped with a digital camera (Nikon, Japan).

### Immunofluorescence and Histological analysis

Mice were subjected to transcardiac perfusion with PBS followed by 4% paraformaldehyde (PFA) after anesthesia with 10% chloral hydrate. Brains were dissected and fixed with 4% PFA overnight at 4^°^C. Tissues were embedded in paraffin after dehydration with graded concentrations of ethanol and xylene or in OCT after cryoprotection with 30% sucrose in PBS. Paraffin-embedded brains were sectioned coronally in 8 μm or 12 μm thick using a microtome (MICROM HM 325, Thermo Scientific, China). Hematoxylin-eosin staining was done according to standard protocols and then slides were mounted with a mounting medium (Wuxi Jiangyuan, Wuxi, China). Images of brain sections were taken using a stereomicroscope (Nikon, Japan) equipped a camera (ToupTek, China) to obtain the whole view of desired ventricular compartments or using Nikon eclipse Ci (Nikon, Japan) equipped with a Micropublisher 6 camera (Teledyne Photometric, Tucson, USA) for imaging lateral ventricles only.

The immunofluorescence on brain sections was done according to standard protocols. Sections were incubated with primary antibodies GT335 (AG-20B-0020, Adipogen, USA, 1:500), mouse anti-acetylated-tubulin (T6793, Sigma, USA, 1:500), mouse anti-Foxj1(eBioscience, France, clone 2A5, 1:200), rabbit anti-Arl13b (17711–1-AP, Proteintech, China, 1:500), rabbit anti-RFP (ab62341, Abcam, USA, 1:500), rabbit anti-vimentin (10366-1-AP, Proteintech, China, 1:200), rabbit anti-Tdtomato (600-401, Rockland, USA, 1:200), rabbit anti-s100b (EP1576Y, Abcam, USA, 1:200) at 4°C overnight and then the immunosignals were visualized by incubation with secondary antibodies Alexa Fluor® 594 donkey anti-rabbit IgG (1:750) and/or Alexa Fluor® 488 goat anti-mouse IgG (1:750) for 1 h at room temperature. For GT335 and acetylated-tubulin co-staining, the IgG isoform specific secondary antibodies anti-mouse IgG2b Alexa Fluor 488 (sms2bAF488-1, Proteintech) and anti-mouse IgG1, Alexa Fluor 568 (sms1AF568-1, Proteintech) were used. Sections were dipped in PBS containing DAPI for 5 min followed by 3 washes with PBS before mounted with antifade mounting reagent (Fluoromount-G (0100-35, SounthernBiotech, USA). To quantify the number of cells positive for specific markers, 3 serious slides with at least 3 sections on each slide were manually counted under immunofluorescence microscope.

### Fluorescence microscopy imaging

All the samples were observed at room temperature under a fluorescence microscope (ECLIPSE 80i; Nikon, Tokyo, Japan) equipped with a 40 × 0.75 NA objective lens (Nikon) or a confocal microscope (TCS SP8; Leica, Wetzlar, Germany) equipped with 63 ×1.4 NA objective lens (Leica). Images were acquired using NIS-Elements software (Nikon) or Las X software (Leica). Image processing was performed using ImageJ and Photoshop (Adobe, California, USA).

### Quantification of the length of multicilia

The length of multicilia was manually measured on images of brain sections stained with Ar13b, GT335, and anti-acetylated tubulin antibody. Stained sections were imaged under a confocal microscope (TCS SP8; Leica) equipped with 63 × 1.4 NA objective lens (Leica) and manually measured on Image J software using segmented line tool to quantify the longest cilia of each multicilia bundle.

### Scanning electron microscopy

Mouse LV walls or tracheas were dissected and fixed overnight at 4°C in 2.5% glutaraldehyde in PBS. After post-fixed in 1% osmium tetroxide, the samples were dehydrated through a series of graded ethanol (30%, 50%, 75%, 95%, and 100%) and critical-point dried using CO_2_ as the transitional fluid. Samples were mounted and sputter coated with gold, and images were captured with a Leo 1530 FEG scanning electron microscope (Zeiss, Germany) at an accelerating voltage of 5 KV.

### Immunoblotting

Mouse tissues were homogenized in the RIPA buffer containing the cOmplete® EDTA-free protease inhibitors (Roche Diagnostics, Mannheim, Germany). After centrifuge at 12,000 rpm for 10 min at 4°C, proteins in the supernatant were separated with 10% SDS-PAGE and then transferred to nitrocellulose membranes. Membranes were blocked with 5% no-fat milk for 30 min, followed by incubation with antibody GT335 (Adipogen, USA, 1:5000) or EP1332Y (Abcam, USA, 1:6000) overnight at 4°C. After washing with 0.1% TBST for 3 times, membranes were incubated with the HRP-conjugated goat anti-mouse or donkey anti-rabbit secondary antibodies (Bioss, Beijing, China) for 2 hr at room temperature. After 3 times wash with 0.1% TBST, the immunoreactivities of proteins were visualized with Western bright ECL reagent (Advansta, Menlo Park, USA).

### Preparation of whole-mounts lateral ventricles

Whole-mounts of the lateral ventricles were prepared as described (Ohata et al., 2014; Pan et al., 2023). Briefly, mice were euthanized with CO_2_, and the lateral ventricles were carefully dissected at a thickness of 500-1,000 μm using Vannas Scissors (66VT, 54140B) in pre-warmed (37°C) dissection solution (25□mM Hepes,117.2□mM NaCl, 26.1□mM NaHCO_3_, 5.3 mM KCl, 1.8 mM CaCl_2_, 0.81□mM MgSO_4_, 1□mM NaH_2_PO_4_·2H_2_O, and 5.6□mM Glucose, pH 7.4).

### High-speed imaging of ciliary beating and whole-mount immunostaining

Live imaging of ciliary beating and immunostaining were performed as described (Ohata et al., 2014; Pan et al., 2023; Zhao et al., 2021) with minor modifications. For high-speed imaging of ciliary beating, the whole-mount preparations were incubated with SiR-tubulin (100 nM; Spirochrome, SC002) in DMEM (Thermo Fisher, 12430062) supplemented with 0.3 mg/ml glutamine, 100 U/ml penicillin, and 100 U/ml streptomycin for 1 h at 37℃ to label the cilia. Ciliary beating was captured with 5 ms exposure time at 100 frames per second (fps) at 37℃ using an Olympus Xplore SpinSR 10 microscope equipped with UPLAPO OHR 60 × /1.50 Objective Lens, Hamamatsu ORCA-Fusion camera, 4,000 rpm CSU Disk Speed, and OBIS Laser.

For immunostaining, freshly dissected whole-mounts of the lateral ventricles were pre-permeabilized with 0.5% Triton X-100 in PBS for 30 sec before fixation to remove soluble proteins and fixed for 15 min in 4% paraformaldehyde. After fixation, the whole-mounts were extracted with 0.5% Triton X-100 in PBS for 15 min and blocked in 4% BSA/TBST blocking solution at room temperature (RT) for 1 h. Next, they were incubated with primary antibodies overnight at 4°C. After three washes with the blocking solution for 5 min each, the whole-mounts were incubated with secondary antibodies at RT for 1h, and mounted with ProLongTM (Thermo Fisher, 2273640). Primary antibodies include mouse IgG anti-acetylated-tubulin (1:1,000; Sigma-Aldrich, T6793), rabbit anti-Centrin 1 (1:200, Proteintech, 12794-1-AP), rabbit anti-β-catenin (1:200, Proteintech, 51067-2-AP), rabbit IgG anti-Cep164 (1:200; Proteintech Group Inc, 22227-1-AP), rat anti-tyrosinated tubulin (1:500; Abcam, ab6160). Secondary antibodies are Alexa Fluor 647 goat anti-mouse IgG (1:1,000; Life Technologies, A-21236), Alexa Fluor 488 donkey anti-rabbit IgG (1:1,000; Life Technologies, A-21206), and Alexa Fluor 568 goat anti-rabbit IgG (1:750; Invitrogen, A-11011). Actin is stained with Phalloidin-TRITC (1:1,000; Sigma, P1951). The confocal images were acquired using Leica SP8 or the Olympus Xplore SpinSR 10 microscope. Three-dimensional structured illumination microscopy (3D-SIM) super-resolution images were acquired using the DeltaVision OMX SR imaging system (GE Healthcare). The z-axis scanning step was 0.125 μm and raw images were processed in SoftWoRx 7.0 software by the following procedures: OMX SI Reconstruction, OMX Image Registration, and maximum intensity projection.

### Quantification of the intensity of actin network around BBs

Quantification of actin enrichment at basal body (BB) patches was performed using Fiji or ImageJ. BB density was estimated using Cep164 intensity. For each cell, BB patches were manually outlined using the polygon selection tool, and the corresponding region of interest (ROI) area was recorded. Within each ROI, the integrated density was measured for both the actin and Cep164 channels. Normalized actin intensity (actin per BB) was calculated as total actin intensity divided by that of Cep164 signals.

### Quantification of cilia beating direction and frequency

The beating direction and frequency were measured using the software FIJI (NIH, USA). For the frequency analysis, the Kymography tool was applied. The R package ggplot2 rendered the analysis of beating directions. To calculate the mean vector length, a previously developed R program was applied (Pan et al., 2024).

### Quantification of BB patch displacement and angle distribution

The BB patch displacement was measured as the ratio of the distance between the center of BB patch and the cell centroid to the distance between the cell centroid and cell boundary in the same direction as described (Mirzadeh et al., 2010; Ohata et al., 2014). These distances and cell centroids are determined using Fiji software. The statistics was analyzed using Student’s *t* test.

To determine the BB patch angle distribution in a given field, vectors were drawn from the cell centroids to the centers of BB patches. The angles of the vectors were measured in Fiji software. Deviations of individual cell’s BB patch angle from the median within the field were calculated and plotted as a histogram using a program that we generated using MATLAB. The percentages were calculated with a bin size of 10°. Ependymal cells within 5 different fields (120 μm x 120 μm for each field) were analyzed. The statistics for the BB angle distribution was analyzed using Watson’s two-sample U^2^ test according to (Zar, 1999) in MATLAB.

## Statistical Analysis

Data are presented as the mean ± SEM and the statistics was analyzed using Student’s *t* test unless otherwise specified. Kaplan-Meier Curves between the wild-type and *Agbl5^M1/M1^*mice was proceeded with a log rank (Mantel-Cox) test. All statistical analysis was performed with GraphPad Prism, p<0.05 were considered statistically significant.

## Data Availability Statement

The lists of assessed predicted off-targets for the mutant mice were provided in Table S1 and S2. Other original data and program codes are available upon request.

## Acknowledgements

We thank Dr. Corey Powell from Consulting for Statistics, Computing, and Analytics Research (CSCAR), University of Michigan for assistance with ciliary beating direction analysis, Dr. Eric Rentchler from the Microscopy Core of Biomedical Research Core Facilities, University of Michigan Medical School for advances on ciliary beating frequency analysis. This work is partially supported by the start-up to HYW from Tianjin University and Elizabeth E. Kennedy Children’s Research Award to HYW from the Department of Pediatrics, University of Michigan Medical School.

