## Supplemental Fig S1-S9 for "Cytosolic Carboxypeptidase 5 maintains mammalian ependymal multicilia to ensure proper homeostasis and functions of the brain"

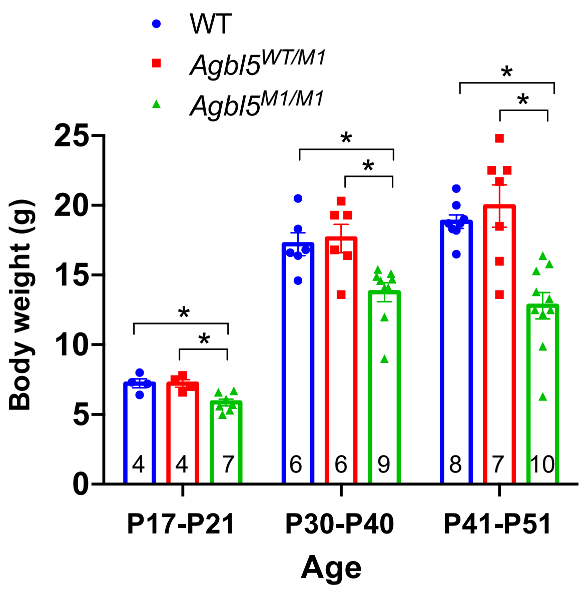


**Figure S1. The growth of *Agbl5*$^{M1/M1}$ mice retards during development.** Compared with the wild-type litter mates, *Agbl5*$^{M1/M1}$ mice showed reduced body weight between the 2$^{nd}$ and 3$^{rd}$ week, and hardly grew after P30 (*, $p<0.5$, student *t* test).

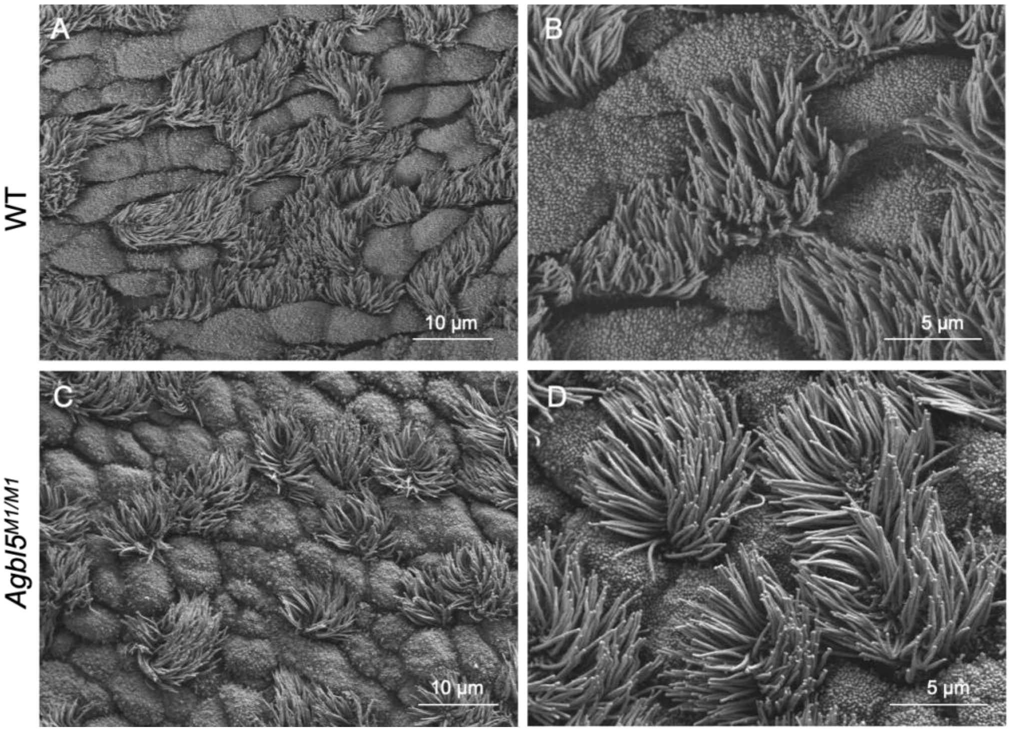


**Figure S2. Scanning electron microscopy analysis of tracheas of *Agbl5*$^{M1/M1}$.** The multicilia of tracheal epithelium in P30 wild-type mice (A, B) point to the same directions, those in the age-matched *Agbl5*$^{M1/M1}$ (C, D) littermates radiated to different directions in the same cluster. Scale bar: 10 μm for A, C; 5 μm for B, D.

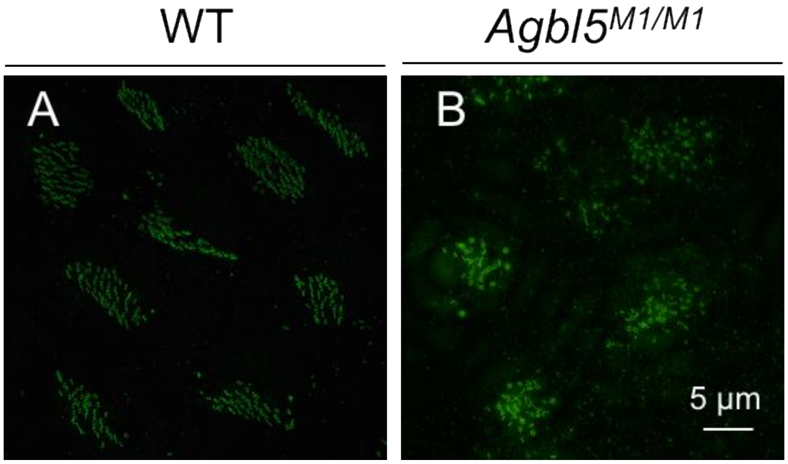


**Figure S3. SIM images of basal bodies in wild-type and *Agbl5$^{M1/M1}$*ependyma.** While basal bodies in wild-type (WT) ependyma are well aligned in individual clusters (A), those in the mutant are not organized (B). Scale bar, 5 μm.

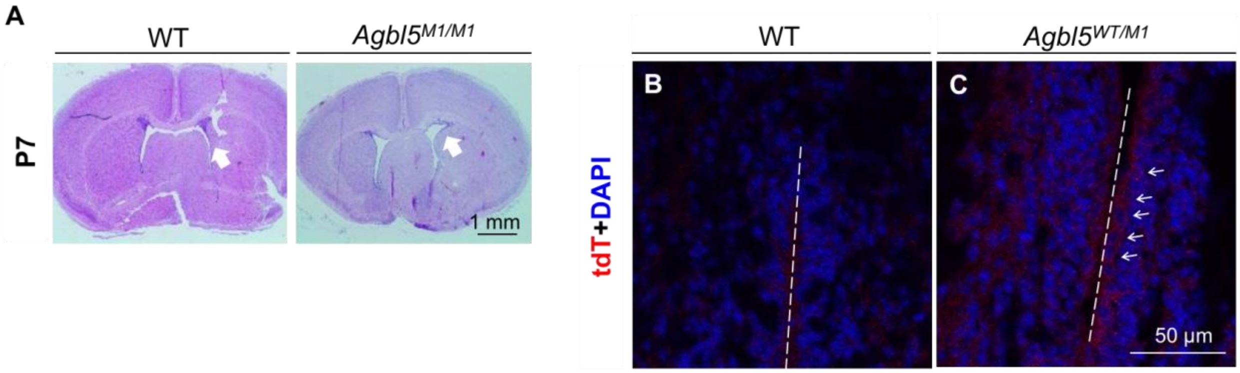


**Figure S4. Expression of *Agbl5* in ependymal cells at early postnatal stage.** (A) The enlarged lateral ventricles in *Agbl5*$^{M1/M1}$ mice were first observed at the age of P7. (B-C) Brain sections of P7 wild-type (WT, B) and *Agbl5*$^{WT/M1}$ (C) mice that were immunostained with tdTomato antibody revealed the expression of *Agbl5* in developing ependymal cells (arrows). Scale bar, A, 1 mm; B, C, 50 µm.

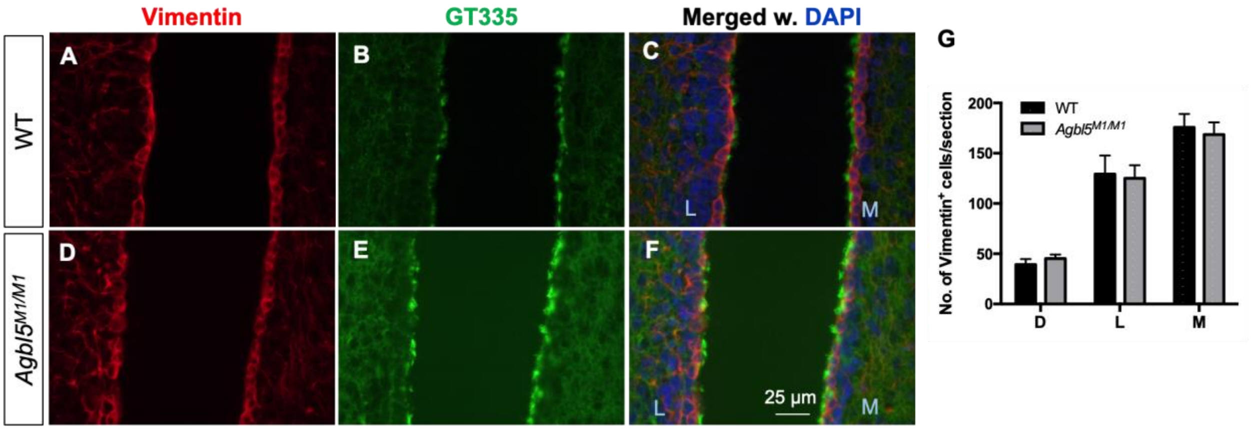


**Figure S5. The commitment of ependymal cells was not affected in *Agbl5*$^{M1/M1}$ mice.** (A-F) Lateral ventricles from P7 wild-type (A-C) or *Agbl5*$^{M1/M1}$ (D-F) mice were co-immunostained with vimentin and GT335 with nuclei stained with DAPI. (G) Quantification showed that the number of vimentin positive cells in individual walls of LV is comparable between wild-type (n=3) and mutant (n=5) mice. L, Lateral wall; M, middle wall; Scale bar, 25 μm.

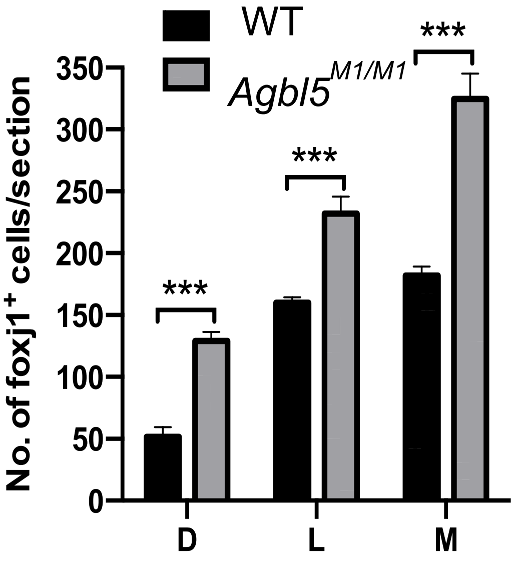


**Figure S6. Number of total Foxj1$^+$ cells in each wall of lateral ventricles per section of P7 mice.**

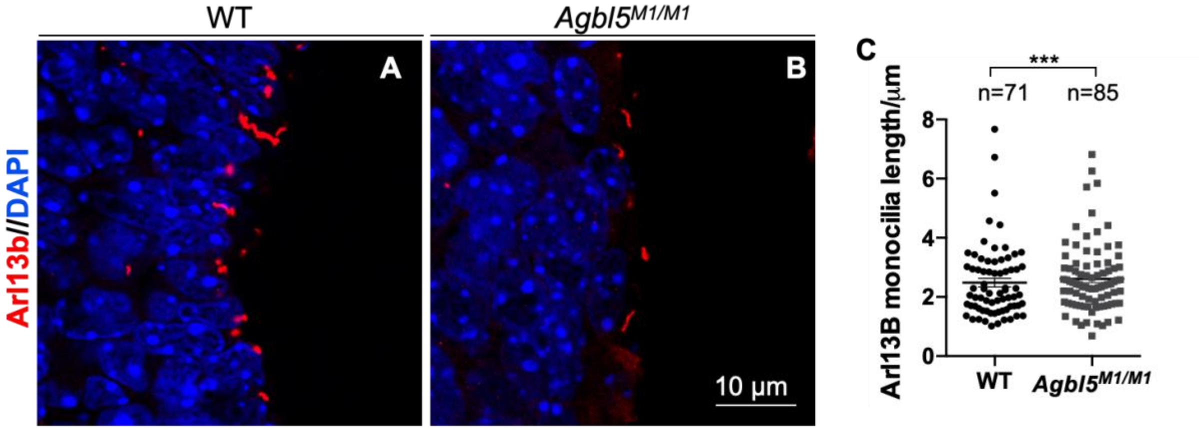


**Figure S7. The length of primary cilia in differentiating ependymal cells is not changed in *Agbl5*$^{M1/M1}$ mice.** (A) Lateral ventricles from P7 wild-type (A) or *Agbl5*$^{M1/M1}$ (B) mice were immunostained for Arl13b with nuclei stained with DAPI. (C) Quantification showed that the length of Arl13b signals in primary cilia is comparable between wild-type (n=4) and mutant (n=3) mice. Scale bar, 10 µm.

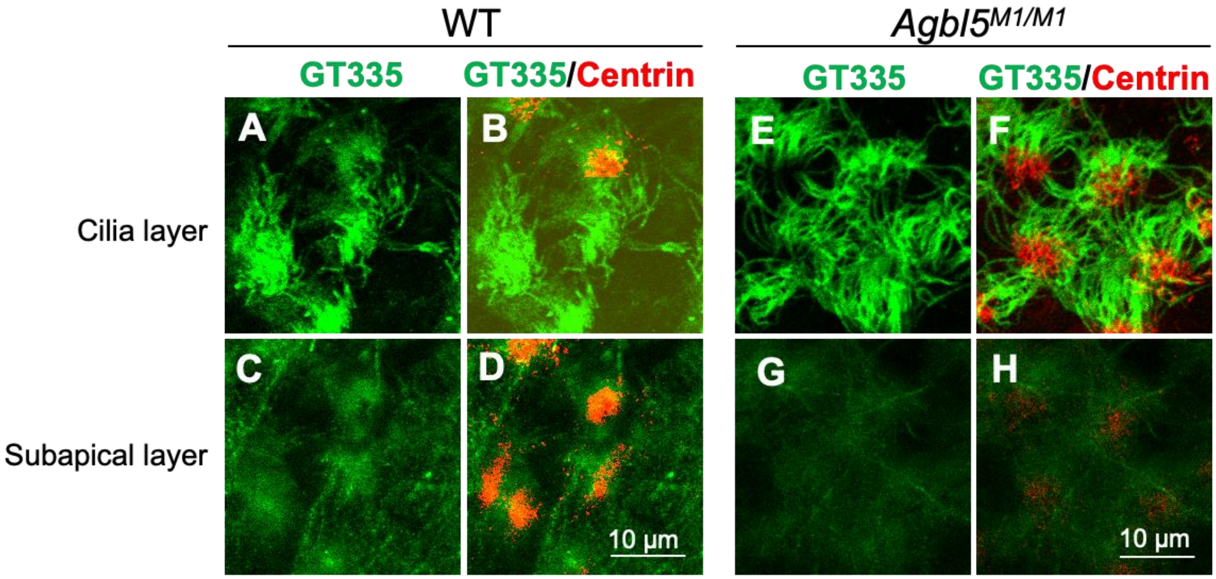


**Figure S8. The glutamylation signals are not increased in the subapical layer in *Agbl5*$^{M1/M1}$ mice.** Whole-mount lateral wall of LVs from P17 wild-type (A-D) or *Agbl5*$^{M1/M1}$ (E-H) mice were co-immunostained for glutamylation (GT335) and Centrin for BBs. While length of GT335 signals is increased in the multicilia of the mutant ependyma (E, F) compared to that in the wild-type (A, B), the GT335 signals beneath the BBs were not increased in the mutant (G, H). Scale bar, 10 μm.

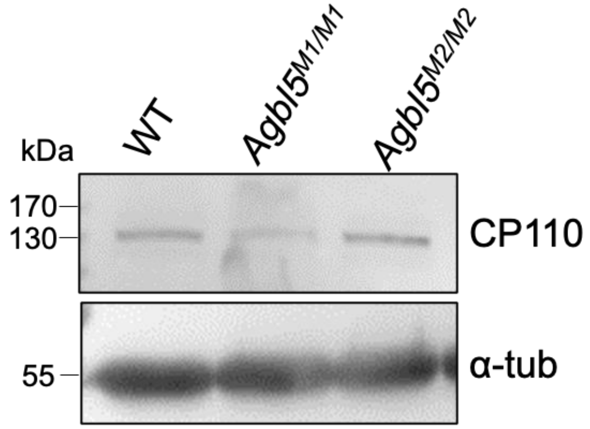


**Figure S9. Immunoblotting analysis of the level of CP110 in ventricles of different genotypes.**

**Movie 1**
Representative high-speed imaging of ependymal cilia from P45 wild-type mice showed their directional beating pattern.

**Movie 2**
Representative high-speed imaging of ependymal cilia from P45 *Agbl5*$^{M1/M1}$ mice for the remnant cilia that hardly beat.

**Movie 3**
Representative high-speed imaging of ependymal cilia from P45 *Agbl5*$^{M1/M1}$ mice for the remnant cilia that beat asynchronously.

**Movie 4**
Representative high-speed imaging of ependymal cilia from P15 wild-type mice showed the unidirectional beating pattern among cells.

**Movie 5**
Representative high-speed imaging of ependymal cilia from P15 *Agbl5*$^{M1/M1}$ mice showed that the cilia of neighboring cells often beat in different directions. Arrows point to cilia in the same cluster that move in different directions.
