## Supplemental Table 1-3 for "Cytosolic Carboxypeptidase 5 maintains mammalian ependymal multicilia to ensure proper homeostasis and functions of the brain"

**Table S1. Identification of off-targets for sgRNAs used in generation of the *Agbl5^M1^* allele via CRISPR/CAS9***

| **Upstream sgRNA:** | | | | |
| --- | --- | --- | --- | --- |
| **Chromosome position** | **MM** | **Gene name** | **Forward primer sequence** | **Reverse primer sequence** |
| chr6:55499397-55499419 | 3 | *Adcyap1r1* | CCTTCAGGTACATCTGCTCACA | TCTACACCAGGCAGGGACAG |
| chr13:60607541-60607563 | 4 | *Dapk1* | CATGAGGCTGGTGGTATTTAT | GCATGTCAACACAACTGACC |
| chr1:184046844-184046866 | 4 | *Dusp10* | CAGGAAAGCCCAGGGGT | TAAATAGCAATGTGATCAGCCTGAA |
| chr11:5717586-5717608 | 3 | *Urgcp* | AAAACATCCAGGTCCTCCTT | CGAGAAGACAGTGTGGTCCT |
| chr6:100774905-100774927 | 4 | *Gxylt2* | GGACTGGAATGATGGCTTAG | CTTCATCATTGCTTTTAGGAAA |
| chr9:50659298-50659320 | 4 | *Dlat* | TGGGTCACTCACCTTCTGAT | GTAATGTGGCGCGTCTGT |
| chr4:117838144-117838166 | 4 | *Slc6a9* | CAGGAAGCAGCCAGGTGG | GGACACACAGGTTCCAGCAA |
| chr14:14756011-14756033 | 4 | *Slc4a7* | GTGGAGCAGTCAAGCCATTGTC | CTGATAATTGAATACAGAGGCAGGAG |
| chr9:78443288-78443310 | 3 | *Mb21d1* | AAATATATGGCGGGAACGTA | GGCAAATCTAGCCATCTGAG |
| chr2:166354157-166354179 | 4 | *Gm14268* | GCTCCTCTTCTTCATTCAGC | AGGAAACAAAGCCATCTGAG |
| chr19:52017492-52017514 | 4 | *-* | TGGAATATCCCTGCCACCTTAGTTC | TTTCTCTGCGTTGCTTTCTGTAGCAT |
| **Downstream sgRNA:** | | | | |
| **Chromosome position** | **MM** | **Gene name** | **Forward primer sequence** | **Reverse primer sequence** |
| chr15:73650582-73650604 | 4 | *1700010B13Rik* | CAAACTCTCCATGAAGCACA | AGTCCAGGTACAGAGCCTTG |
| chr11:69653088-69653110 | 4 | *Fxr2* | CCACCCAGATTTGACAGTTC | GGTTATTTCCTCCCAACTCC |
| chr11:66488144-66488166 | 4 | *Shisa6* | GAGCCATGTTTGTGCTGAT | ATGCGTGAATCTCATTTGTG |
| chr3:74040980-74041002 | 4 | *Gm20356* | CCTCTTTGGTCCCATTTTC | GCCATGAACCTTTCAGAGAC |
| chrX:105870181-105870203 | 4 | *Atrx* | TGGAGAGGACCTGAGCTTTA | TTTCTCTTTCCTTGGTTTGG |
| chr7:102205970-102205992 | 3 | *Nup98* | TGTTCCAGGGCCAGTGAGAT | CCTGGCATGGAGGCTAGGAA |
| chr6:55037581-55037603 | 4 | *Gars* | CCCACTGGAACACCCATACT | ATTAACTGGCATGCCCTGAG |
| chr5:142769309-142769331 | 4 | *Tnrc18* | AGAGAACGAGAACGAACCTTCCAG | AGACAGATATGAAGCCCAGGCTG |
| chrX:33287193-33287215 | 4 | *Gm15274* | ACAAGGCAGCCCAGCTAG | GATGTTGGCTATTGGCTTCC |
| chrX:32947754-32947776 | 4 | *Gm21789* | ACATCCTGATTACAGCCTCTCCTCC | TGGTACATCCTCTCCCACTGAGG |
| chr12:6447103-6447125 | 4 | *NA* | AGCTAGCCAGTGGAACTATCGC | CCAAAGGCAAATGATGTTCTCAGAATTCA |

***** Names and genomic location of genes that were predicted as off-targets with maximal four bases mismatches. MM: Mismatch.

**Table S2. Identification of off-targets for sgRNAs used in generation of the *Agbl5^M2^* allele via CRISPR-CAS9***

| **Upstream sgRNA** | | | | |
| --- | --- | --- | --- | --- |
| **Chromosome position** | **MM** | **Gene name** | **Forward primer sequence** | **Reverse primer sequence** |
| Chr11:50022372-50022394 | 3 | *Intergenic* | TGGCAGAGCAATTGTCCTGCATG | CCTACTCACTGGTGTCTGGTTTC |
| Chr17:29724904-29724926 | 3 | *Intergenic* | GAGCCATTACATTGGCTATGCCAG | CTTTCCGAAGGCTGCTCAGTG |
| Chr14:54570699-54570721 | 3 | *ajuba-intron4* | TTCCCGAGACAGGGTTTCTCTGTA | CGTTCCTTTCCACCACCAACC |
| Chr13:43377282-43377304 | 3 | *sirt5-intron4* | TCCCTGCTGACGACAAGGGAT | CGGGGCATACCACAAGAAACC |
| Chr11:61119960-61119982 | 3 | *Intergenic* | CACAGACATCCCTGGGATGG | CCCTCAGTGCCTTTGATGATGAAC |
| Chr17:15669812-15669833 | 2 | *Intergenic* | GGGAGTATTGGTGTGAAAGGCC | AGGACAGCCAGGGCTACACA |
| Chr13:48047126-48047147 | 2 | *A330033J07Rik-intron2* | CTGTCATTTCCAACAGAACACCCACTTTAG | GCAAGCTGGGAATTTCCTTTGCTG |
| Chr1:192859610-192859631 | 2 | *Intergenic* | GGAAGCACTCCTGGTACCTTTTCT | GACAAGCCAAGAGGAGATATGAGG |
| Chr13:54496178-54496199 | 2 | *Intergenic* | AGCCAAATAAACCCTTTCCTCCCC | AAATCCACCTGCCTCTGCCTC |
| Chr15:48299282-48299303 | 2 | *csmd3-intron6* | GGTGAGACCAATTGTTGTTGTTGTTGAAGG | TTTGGAAAGGGGCAGAAAGGTGCA |
| Chr11:88557902-88557923 | 2 | *msi2-intron6* | CCTGCCTAGTTTCACCCTAGC | GCGAGTTTATGAAAGGGACCCAG |
| Chr4:148579132-148579153 | 2 | *Intergenic* | GTCATGTCAGACCCTAGACTCAC | CCTGTCCCCTACAGATTCTAACC |
| Chr13:48047124-48047147 | 2 | *A330033J07Rik-intron1* | CTGTCATTTCCAACAGAACACCCACTTTAG | GCAAGCTGGGAATTTCCTTTGCTG |
| Chr12:90515027-90515050 | 2 | *NA* | CTCTGTATAGTCTTGGCTGTCCTG | AGAGTCAGAGTCTAACAGAGGGTG |
| **Downstream sgRNA** | | | | |
| **Chromosome position** | **MM** | **Gene name** | **Forward primer sequence** | **Reverse primer sequence** |
| Chr7:139945453-139945475 | 3 | *Intergenic* | TGAAAGTTCTAGTTGGGCAGCTGG | CTGGCTGGATAAGGTACGTGG |
| Chr4:127232539-127232560 | 2 | *Dlgap3-intron8* | GGGGCCTGAACGTTTCTTGTC | GGTAGACAAAGACCCATGGACCT |
| Chr4:128756772-128756793 | 2 | *A3galt2-intron1* | GGGAGGTTTGGTGAAGGTAGC | GCTGTTTATACACGGAAGCTCCTC |
| Chr7:139945452-139945475 | 2 | *NA* | TGAAAGTTCTAGTTGGGCAGCTGG | CTGGCTGGATAAGGTACGTGG |
| chr9:112951190-112951212 | 4 | *NA* | TCTCATTGAAAACTCAAGAATAACA | ACAAGTATGCTGCTGTGTGC |
| chr11:116533798-116533820 | 4 | *Sphk1* | ATGCCAGTTCTGGGAATTGT | AAAGAGTGGTGGGGTTCTCT |
| chr5:31323504-31323526 | 4 | *Gckr* | AGGTGCTTGCTACCAAATCTT | CTCTGAAGGCAGAAGTGGTG |
| chr4:94962199-94962221 | 4 | *Mysm1* | TGTGTTACAAGTATTCTTTCCCCA | GCAAAGCTGGGAAAACCTTG |
| chr10:97314596-97314618 | 4 | *NA* | GTGACAAGGCATCAAAACACA | AGAACCAGGGGAGATTCTTAAC |
| chr5:129737441-129737463 | 4 | *Gbas* | ACGCTCGGCTAATTTTTAAG | CAGCTCCAGGGGACCTAATA |
| chr6:117985065-117985087 | 4 | *Gm4875* | AGCCATGACCTTTCAAACACT | TGCCCGAAAAGCATTAAGCA |
| chr6:8002193-8002215 | 4 | *Col28a1* | AGCTGAGTCTGTGAGGAAGG | TTTCCACTGTCCACTCACCT |

***** Names and genomic location of genes that were predicted as off-targets with maximal four bases mismatches. MM: Mismatch.

**Table S3. Primers used for genotyping different *Agbl5* alleles**

| **Strains** | **Allele** | **Genotyping primers** | **Predicted sizes of PCR products (bp)** |
| --- | --- | --- | --- |
| ***Agbl5^M1^*** | wild-type | F: 5’-TCTGCCCTTCCTCCCTATCC-3’  R: 5’-CACCCCCTGCAGTCCTCATA-3’ | 327 |
|  | mutant | F: 5’-TCTGCCCTTCCTCCCTATCC-3’  R: 5’-TGTAGATCAGCGTGCCGTCC-3’ | 455 |
| ***Agbl5^M2^*** | wild-type | F: 5’- TAGAGCTGGAGGAGTGACAG-3’  R: 5’-CCATGGCAGGAAGTGATTGT-3’ | 359 |
|  | mutant | F: 5’- TAGAGCTGGAGGAGTGACAG-3’  R: 5’-CGGCAATATGGTGGAAAATA-3’ | 571 |
